# The supramolecular organization of SARS-CoV and SARS-CoV-2 virions revealed by coarse-grained models of intact virus envelopes

**DOI:** 10.1101/2021.09.16.460716

**Authors:** Beibei Wang, Changqing Zhong, D. Peter Tieleman

## Abstract

The coronavirus disease 19 (COVID-19) pandemic is causing a global health crisis and has already caused a devastating societal and economic burden. The pathogen, severe acute respiratory syndrome coronavirus 2 (SARS-CoV-2), has a high sequence and architecture identity with SARS-CoV, but far more people have been infected by SARS-CoV-2. Here, combining structural data from cryo-EM and structure prediction, we constructed bottom-up Martini coarse-grained models of intact SARS-CoV and SARS-CoV-2 envelopes. Microsecond molecular dynamics simulations were performed, allowing us to explore their dynamics and supramolecular organization. Both SARS-CoV and SARS-CoV-2 envelopes present a spherical morphology with structural proteins forming multiple string-like islands in the membrane and clusters between heads of spike proteins. Critical differences between the SARS-CoV and SARS-CoV-2 envelopes are the interaction pattern between spike proteins and the flexibility of spike proteins. Our models provide structural and dynamic insights in the SARS virus envelopes, and could be used for further investigation, such as drug design, and fusion and fission processes.

## Introduction

Coronaviruses, including severe acute respiratory syndrome coronavirus (SARS-CoV) and currently in particular SARS-CoV-2 are a major threat to public health (1). They are enveloped positive sense RNA viruses that can be transmitted from animals to humans and cause a variety of diseases ranging from common cold to severe diseases (2).

The SARS-CoV and SARS-CoV-2 virions contain four main structural proteins: the nucleocapsid (N), spike (S), envelope (E) and membrane (M) proteins (3). Ns are tightly packed with RNA in the viral lumen, while S, M and E proteins are located on the lipid bilayer of the viral envelope. E (about 75 amino acids) is a small hydrophobic integral membrane protein, and a multifunctional protein, supposed to be involved in virus assembly and release, and pathogenesis (4-6). M (about 220 amino acids) is the most abundant structural protein in CoV virions and is composed of three parts: a short N-terminal domain at the virion exterior, three transmembrane (TM) helices and a carboxy-terminal domain at the virion interior (2, 7, 8). M is the primary driver of the virus budding process and directs the virion assembly by interacting with other structural proteins (7, 9, 10). S mediates the fusion process between viral and host membranes (11). It is a homotrimer. Each monomer consists of two subunits: S1 (at the N-terminus, responsible for receptor recognition and binding) and S2 (at the C-terminus, directing the subsequent fusion process) (11, 12). The cryo-EM structures of SARS-CoV-2 S revealed its shared architecture with SARS-CoV S, while the sequence of SARS-CoV-2 S shares about 77% identity with that of SARS-CoV S (13). Both Ss recognize and bind angiotensin-converting enzyme 2 (ACE2), a zinc metallopeptidase involved in cardiovascular and immune systems regulation, to enter and infect human cells (14, 15).

However, the toxicity and transmission capacity of SARS-CoV and SARS-CoV-2 are significantly different. In light of the ongoing global health emergency, there is an urgent need to clarify how the envelope of the CoVs fulfills its function and explore why the infection capacity is different. New structures of SARS-CoV-2 proteins have been obtained by cryo-electron microscopy (cryo-EM) nearly weekly (16-21). Computational approaches have also been used for structural prediction of unresolved sections (22), molecular dockings of different drug molecules on virus proteins (23-26), and free energy calculations of the S-ACE2 binding process (27-32) and the down-up transition of the receptor binding domain (RBD) (33, 34). The larger scale spatio-temporal processes, such as virion assembly, virus architecture, fusion, and budding, are still poorly understood, and remain challenging for experimental techniques as well as all-atom molecular dynamics (MD) simulations (35).

Coarse-grained (CG) models have been proven to be powerful to probe spatio-temporal large-scale process of complicated biomolecular systems (36). Martini CG models have been widely used to investigate protein-lipid interactions (PLIs) and protein-protein interactions (PPIs) (37, 38). In this study, we constructed Martini CG models of SARS-CoV and SARS-CoV-2 envelopes, containing multiple copies of E, M and S proteins and thousands of different lipids; then microsecond (μs) MD simulations were performed to equilibrate the CG models. Overall, our simulations revealed structural and dynamic details of the virus morphology, conformations of three structural proteins (E, M and S), and their PPIs and PLIs for both SARS-CoV and SARS-CoV-2. Our results provide insight into structural and dynamic details of the critical difference between SARS-CoV and SARS-CoV-2 envelopes. Coordinate files of the CG models are available at GitHub (https://github.com/ChangqingZhong/Martini-MD-of-SARS-CoV-and-SARS-CoV-2).

## Results

We constructed Martini CG models of SARS-CoV and SARS-CoV-2 envelopes with Es, Ss and Ms inserted in a lipid vesicle (Fig. 1A). The lipidomics of the vesicle is according to the composition of the human endoplasmic reticulum (ER)-related membrane (39), and is asymmetric between the outer and inner leaflets (Table S1). Microsecond MD simulations were carried out to equilibrate the models. Longer simulations (up to 12 μs) did not result in observable changes in the virions, so some simulations were performed for 3 μs to limit the computational cost (Table S2).

**Figure 1.**
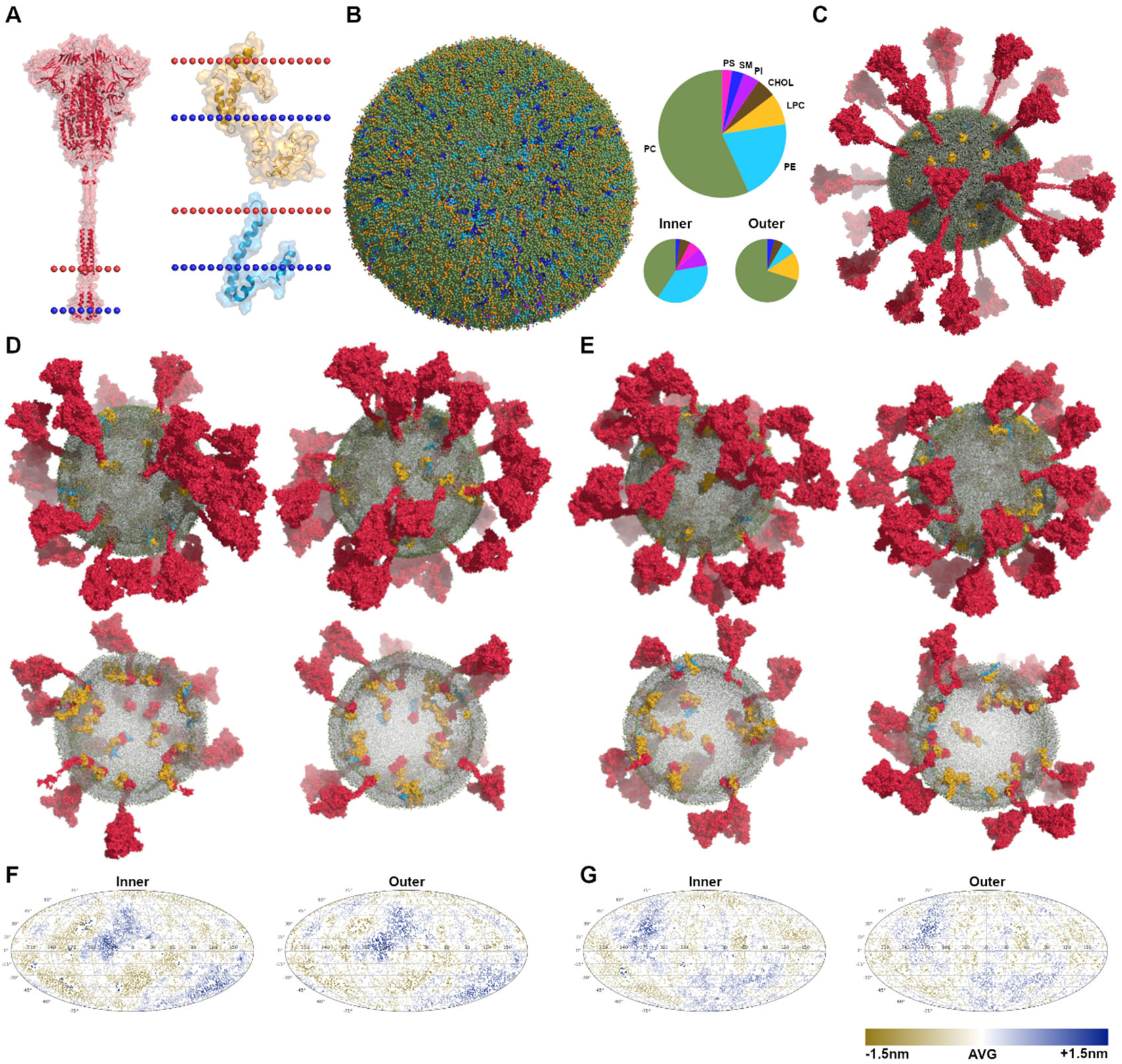
Martini CG models of virus envelopes. (A) The structures of S (in red), M (in yellow) and E (in blue) with the dotted lines indicating the position of lipid bilayers. (B) The equilibrated vesicle with the lipid composition according to that of the endoplasmic reticulum (ER). The lipid bilayer is asymmetric, the compositions of inner and outer leaflets are different. Lipid types considered here are phosphatidylcholine (PC), phosphatidylethanolamine (PE), phosphatidylserine (PS), lyso-PC (LPC), sphingomyelin (SM), phosphatidylinositol (PI), and cholesterol (Chol). (C) Multiple copies of proteins (E×15 in blue, S×30 in red, and M×45 in yellow) were inserted into the equilibrated vesicle, making up the initial CG model of virus envelopes. (D)-(E) The equilibrated models after microsecond MD simulations of SARS-CoV-2-md1 and SARS-CoV-md3, illustrated by the side view and transverse section. (F)-(G) Mollweide projection maps of the envelope membranes of SARS-CoV-2-md1 and SARS-CoV-md3, illustrated using the distances between lipid heads and the vesicle’s center of mass.

### The physical coarse-grained models of the virus envelopes

The virion size varies in different reports (17, 21, 40), in the range of 50-120 nm for the outer diameter. In order to minimize the required computing resources, the Martini CG models constructed here, with an average outer diameter of ∼53 nm, have a total of ∼7.3 million CG beads, containing 90 transmembrane proteins imbedded in a membrane of 16,000 lipid molecules with 68% PC, 7% PE, 16% LPC, 5% CHOL, and 4% SM in the outer leaflet, and 41% PC, 37% PE, 9% PI, 5% PS, 5% CHOL, and 2% SM in the inner leaflet (Fig. 1A-C and Table S1).

There is no clear consensus on the stoichiometric composition of structural proteins of the virus envelopes so far. The reported number of Ss per vision varies in the range of 20-40 (17, 21), so 30 Ss were firstly randomly inserted into the equilibrated vesicle. The prefusion conformation was used for all Ss, because the cryo-EM results suggested that more than 95% of Ss on the virions are in prefusion conformation. Then 15 Es, the minimum abundance, were located randomly, and finally limited by the size of our models, as much as 45 Ms were placed, which may be less than previously inferred (35, 41). A total of 90 structural proteins were included, occupying about 15% of the virion surface and resulting in a 1:180 protein-to-lipid ratio. These structural models enable us to probe the nanoscale organization and dynamics of the SARS-CoV and SARS-CoV-2 envelopes.

In all simulations, both the SARS-CoV and SARS-CoV-2 envelopes are stable and present very similar morphology globally (Figs.1D-E and S4-S11). The shapes of both virus envelopes maintain a relatively regular sphere. The spherical morphology has been observed in several cryo-EM studies (17, 20, 21). The distances between individual lipid heads and the center of mass of the entire virion present slight fluctuations less than 1.5 nm. The landscapes of distance fluctuations of the inner leaflet show almost the same pattern as that of the outer leaflet (Figs. 1 and S4-S11), indicating no significant variation of the bilayer thickness and curvature. The local curvature induced by transmembrane proteins was observed in 500 ns atomistic MD simulations and was conjectured to be the driving force of the deviation from an ideal sphere (42). In our virus models, however, no curvature changes in the membrane were observed in all simulations, which may be the reason that the virions maintain a spherical shape.

The average curvature radius (*R*) of the equilibrated vesicle, calculated with MemCurv software (43), before inserting proteins is about 22.2 nm. After inserting proteins, the vesicle shrinks slightly to about 21.9 nm in radius. During all simulations, except for the simulation SARS-CoV-2-md1, *R* and the radius of gyration (R_g_) of envelopes without the ectodomains of S are almost constant with only slight fluctuations, but R_g_ of intact envelopes decreases from 32.8 nm to ∼29.1 nm, resulting from the swaying motions of Ss at all angles (Fig. S12). In the simulation SARS-CoV-2-md1, a remarkable decrease of 0.3 nm appears in the evolution of *R* at around 1.5 μs, induced by the stalk of one S falling on the membrane artificially (Fig. S12 and S13).

### S-S interactions on the envelopes

Overall, the homotrimer Ss organize into curvilinear strings with varying length (up to 11 Ss) and a small number of five-membered rings spatially through extensive PPIs between the huge heads (S1 domains), but without obvious aggregation (Figs. 1, 2A and S4-S11). The morphology of spatial organization is consistent with recent cryo-EM results (17, 18, 21). The SARS-CoV Ss prefer the conformations of dimer and trimer, while SARS-CoV-2 Ss tend to connect into longer oligomers (Fig. 2B). A global picture of S-S interactions may be obtained from radial distribution functions (RDFs) of Ss around Ss. The difference between SARS-CoV and SARS-CoV-2 Ss is also reflected in the average RDFs (Fig. 2C). RDFs show that both first peaks appear at about 3.3 nm, but the intensity and position of the second peaks are quite different. The average second peak of SARS-CoV Ss at about 13.5 nm are weaker than that of SARS-CoV-2 Ss at about 8.7 nm. The positions of the second peak of each Ss are statistically significantly different between SARS-CoV and SARS-CoV-2 with a p-value of 0.02. These results manifest that SARS-CoV-2 Ss has a higher tendency of aggregation than SARS-CoV Ss.

**Figure 2.**
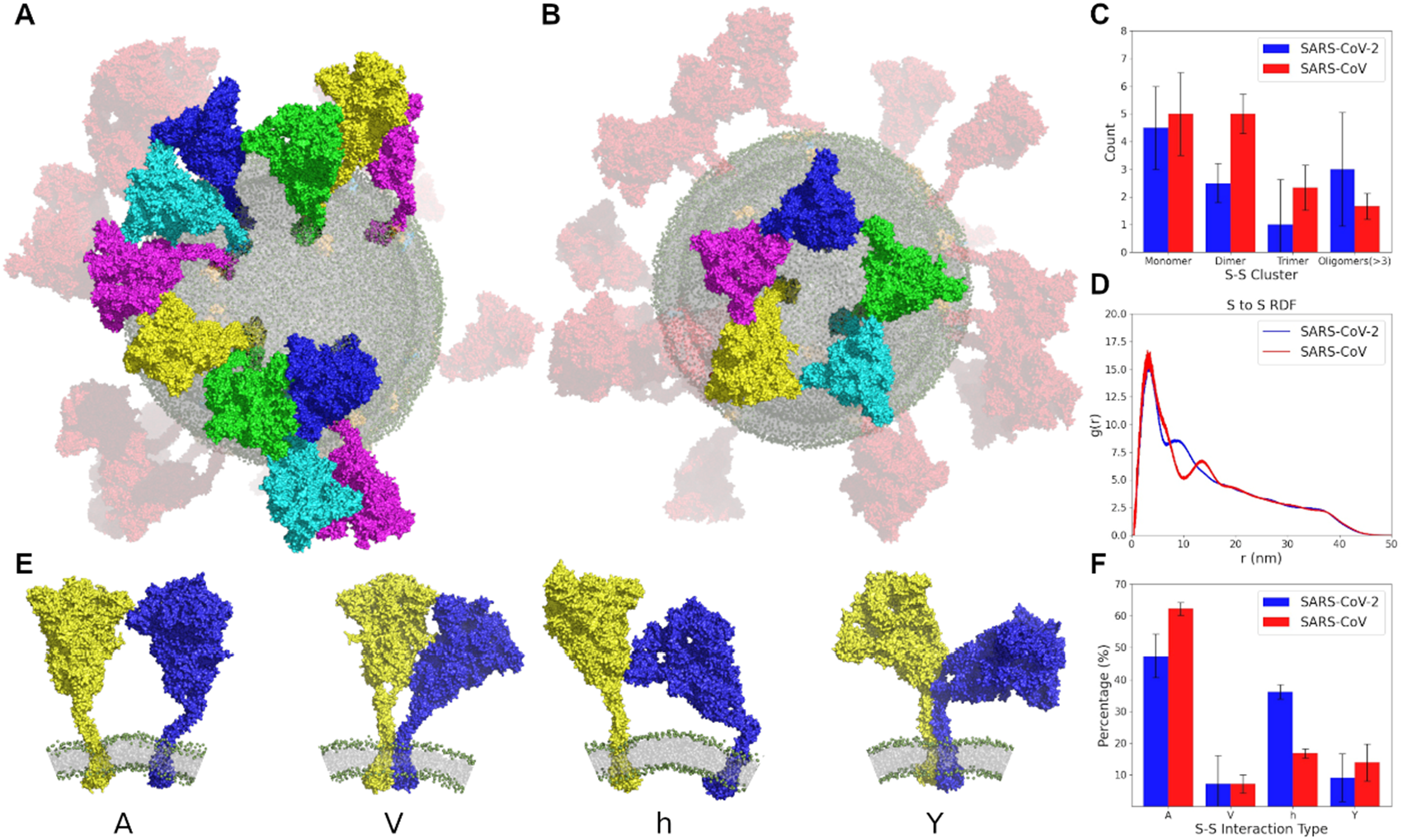
S-S interactions. (A)-(B) Examples of the curvilinear string and five-membered rings spatially organized by S heads. (C) The number of S-S clusters in curvilinear strings with different lengths. (D) The radial distribution function (g(r)) between Ss. (E) The four types of S-S interactions and their percentages (F).

The stalks of Ss in both SARS-CoV and SARS-CoV-2 systems show high flexibility, resulting in orientations in all directions and a variety of PPI patterns. According to interaction sites, the S-S interactions can be classified into four main types: A-shaped, V-shaped, *h*-shaped and Y-shaped (Fig. 2C and 2D). The A-shaped PPI is dominated (about 50% for SARS-CoV-2 and 60% for SARS-CoV) and mainly through the interactions of adjacent N-terminal domains (NTDs), while the V-shaped structure is minor (about 7%), and has extensive interactions. The proportion of the *h*-shaped PPI (NTD-CD) has the greatest difference between SARS-CoV-2 (36%) and SARS-CoV (17%). The Y-shaped PPI contribute about 10% and has intertwining stalks and TMs. Both A-shaped(40) and Y-shaped (21) dimers of S trimers have been reported in recent cryo-EM studies.

### M mediating PPIs on virus envelopes

Visualizing the envelopes without the S ectodomains, it is clear that the TMs of different structural proteins form string-like islands with variable size (up to 18 different proteins) in the envelopes of both SARS-CoV and SARS-CoV-2 (Figs. 3A and S14-S21). The string-like islands was also observed in previous simulations of several different complex membrane models (38, 44). Most of the islands are made up of a mixture of E, M and S proteins. The promiscuous PPIs stabilize the protein islands and will slow their diffusion. The average diffusion coefficients of proteins, calculated from mean-square displacements of the last 1 μs (1.02±0.51 for E, 0.58± 0.28 for S and 0.80±0.42 for M), reduced to about 60% of these calculated from the first 1 μs (1.50±0.42 for E, 1.01±0.58 for S and 1.33±0.34 for M) (Figs. S22). The degree of slowing down is the same as that caused by the crowding of GPCRs (45).

**Figure 3.**
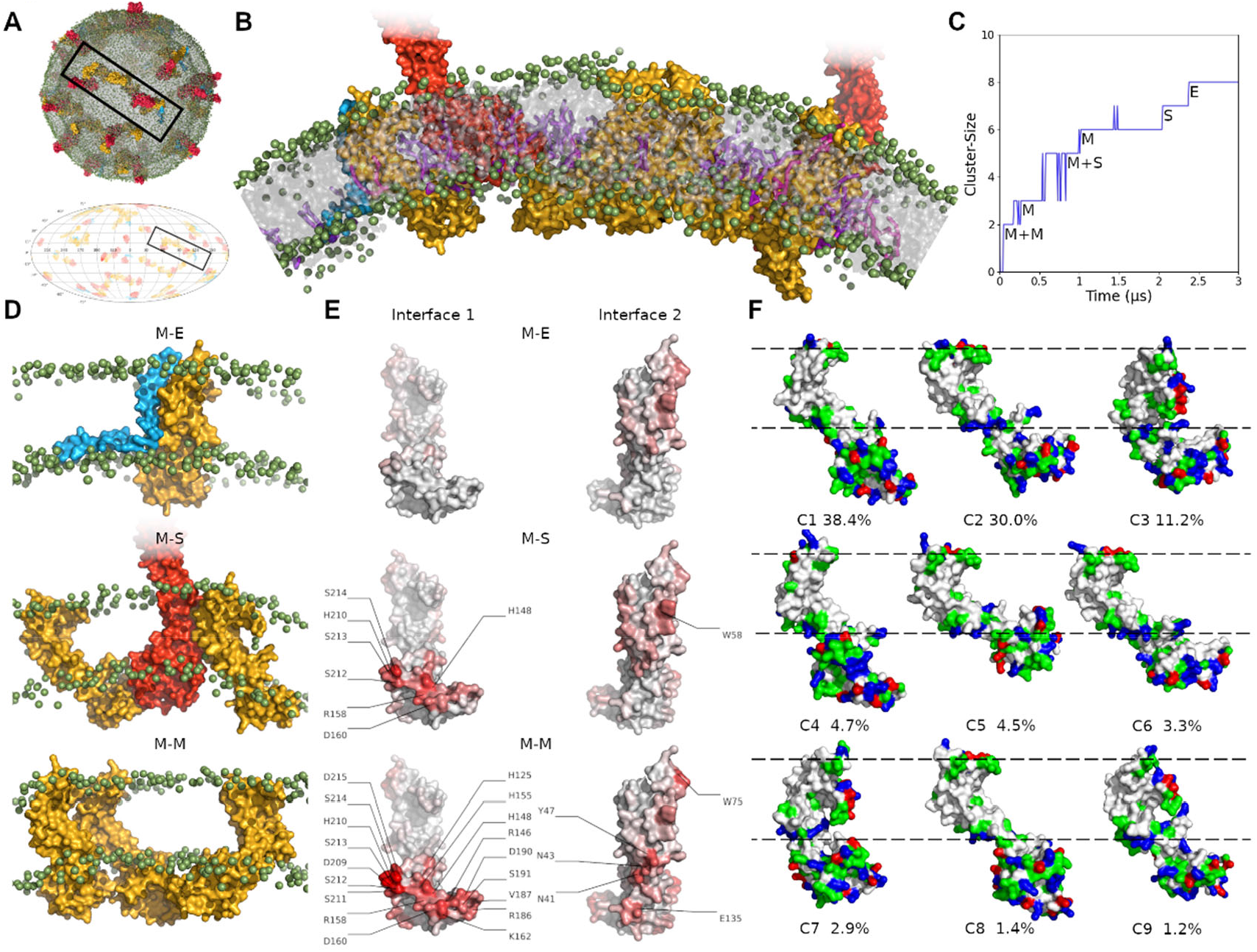
Protein-protein interactions in the envelope membranes. (A) An example of the equilibrated CG model of virus envelopes with S showing only the TM domains, and its Mollweide projection map of protein distributions. The islands in the black frame are enlarged in (B). The color scheme is the same as Fig. 1. The lipid heads are shown in green spheres, and PI lipids in purple sticks. (C) The formation process of this island over the simulation time. (D) Examples of M-E, M-S and M-M interactions. (E) Residues involving in PPIs on the two M interfaces (Interface-1 and Interface-2). The shade of red indicates the contribution. (F) The representative M structure of 9 clusters accounting for more than 1%, resulting from a RMSD-based clustering analysis. The structure was colored according to the physicochemical properties of amino acids: polar residues in green, basic residues in blue, acidic residues in red and nonpolar residues in white.

Es, in monomers in our models, have extensive contacts with Ms and Ss, but no tendency to form pentamers was observed. It is noteworthy that more SARS-CoV Es distribute around M and S than SARS-CoV-2 Es do (Fig. S23).

The recruitment of islands is mainly driven by extensive PPIs between E/M/S and M proteins, the most abundant protein in the virus envelopes. Taking this randomly selected cluster (the one shown in Fig. 3B) as an example, two Ms form a dimer in the first 0.1μs, and then the third M, a M-S cluster, the fourth M, a S and an E were recruited one by one (Fig. 3C). Two preferred interfaces of Ms mediate both homologous and heterologous PPIs (Fig. 3D and 3E). Interface 1 involves the residues located at the ectodomain, while the Interface-2 involves the residues located at the transmembrane domain (Fig. 3E). Promiscuous PPI are largely, but not exclusively, mediated by charged residues displayed on Interface-1, and aromatic residues (W58, W75 and Y47) on Interface-2. Primarily based on these two interfaces, the TM domains form M-S (Interface-1 – S), M-M (Interface-1 – Interface-2), and M-E (Interface-2 – E) PPIs, and assemble into long curvilinear islands. The residues involved in M-S and M-M interactions are almost the same. Previous morphological studies have shown that S protein does not appear in the envelope region in absence of M proteins(41), consistent with our simulations. It indicates that Ms may prevent excessive aggregation of Ss.

Previous morphological studies(41, 46) suggested that a part of Ms form dimers, which also was confirmed in our simulations. The M dimer is tightly bound by the strong and extensive interactions between the two interfaces (Fig. 3D and 3E). In the process of dimer formation, the endodomain domains contact firstly, and then Ms gradually move closer until the TM region have contacts forming the dimer (Fig. S24). The endodomain domain of M mediates the interaction between dimers.

None of previous studies produced a clear picture of the structure of M. In the simulations, in spite of the restriction of elastic networks imposed on proteins, M presents a variety of structures. A root-mean-square-difference-based clustering with a cutoff of 0.5 nm resulted in 24 clusters. The representative structures of 9 clusters accounting for more than 1% are shown in Fig. 3F. The cryo-EM study has classified M structures into elongated and compact conformations according to the length of the endodomain domain (41). Our simulations clearly demonstrated these two conformations. The endodomain has interactions with the lipid heads, resulting in the compact conformation (∼55%, clusters C2, C3, C5, C6, C7 and C9), while the elongated conformation (∼45%, clusters C1, C4 and C8) appears an extended endodomain domain, which may interact with the RNA package (47).

### Lipid micro-environments of structural proteins

Protein modulates its local lipid environment in a unique way and lipid‒protein interactions is regarded as unique fingerprints for membrane proteins (48). We calculated the lipid depletion-enrichment (D-E) index of the three structural proteins separately (Table 1). The D-E index greater than 1 means enrichment and less than 1 means depletion. Consistent with the protein-protein interactions discussed above, the lipid environments of SARS-CoV and SARS-CoV-2 models are consistent due to the highly similar TMs.

**Table 1.** The D‒E index matrix, with average depletion/enrichment for different structural proteins. The D‒E index is computed by dividing the lipid composition of the first 0.7 nm shell by the bulk membrane composition.

| SARS-CoV | PC | PE | Chol | SM | PS | PI | LPC |
| --- | --- | --- | --- | --- | --- | --- | --- |
| S | 0.56±0.02 | 1.67±0.07 | 1.03±0.04 | 0.21±0.03 | 1.72±0.06 | 3.43±0.09 | 1.26±0.03 |
| M | 0.55±0.01 | 1.56±0.04 | 0.88±0.04 | 0.25±0.03 | 2.03±0.07 | 3.92±0.21 | 1.31±0.05 |
| E | 0.57±0.01 | 1.21±0.03 | 2.22±0.34 | 0.51±0.04 | 2.18±0.32 | 4.67±0.21 | 0.81±0.12 |
| SARS-CoV-2 | PC | PE | Chol | SM | PS | PI | LPC |
| S | 0.56±0.01 | 1.72±0.05 | 1.02±0.10 | 0.22±0.01 | 1.65±0.25 | 3.27±0.22 | 1.19±0.04 |
| M | 0.53±0.01 | 1.61±0.02 | 0.83±0.01 | 0.21±0.02 | 1.98±0.12 | 4.03±0.04 | 1.28±0.03 |
| E | 0.59±0.02 | 1.17±0.04 | 2.25±0.09 | 0.46±0.07 | 1.95±0.30 | 4.66±0.23 | 0.77±0.01 |

Compared with M and S, E presents a significantly different lipid microenvironment. The depletion of PC and SM and the enrichment of PE, PS and PI are consistent for all the three proteins. However, it is noteworthy that Es have more SM and CHOL in the first layer of lipids. In the simulations, CHOLs bind non-specifically and dissociate on a sub-microsecond time scale. Protein tends to located in its unique lipid environment. The difference of lipid micro-environments between M/S and E is also reflected in our simulations by most of Es locating at the end or inflection point of the string-like protein islands (Figs 3A, 3B and S14-S21). In other words, the lipid micro-environment of E monomer may a signal to terminate the further extension of the protein clusters and may contribute to the pattern of protein distribution.

PIs, with four negative charges, are highly enriched around all proteins, mediating the protein clusters (Fig. 3B). Positively charged residues are abundant at the layers of lipid heads, and have strong electrostatic interactions with PI heads. These interactions ensure the stable embedding of proteins in the lipid bilayers, especially for S, which has a large and highly flexible head but only three TM helices.

### The flexibility of Ss on the envelopes

Ss show high flexibility with the heads orienting in all different directions (Figs. 1, S4-S11, 4A and 4B). The tilt angle α was defined as the angle between the orientation of the central helix and the normal axis of the envelope bilayer. The α distribution of SARS-CoV-2 Ss ranges from 1.1 º to 116.2 º with a densest population at about 60°, while the α distribution of SARS-CoV Ss shows an almost equally wide range (0.3 º to 100 º) and a densest population at about 35°. The tilt angle distribution of SARS-CoV-2 Ss is consistent with some of the recent cryo-EM results, which also showed a peak at about 60° (17). The distribution of α indicates that SARS-CoV-2 Ss prefer a more standing conformation, comparing with SARS-CoV Ss. The sequences of the stalks are conserved between SARS-CoV and SARS-CoV-2, so the difference in distribution may result from the different PPI patterns. The flexibility of S makes it easier to search for receptor proteins, but excessive flexibility is not conducive to the formation of stable interactions. Therefore, it may be one of the reasons why SARS-CoV-2 has a stronger infection ability.

Two conformations of Ss, “RBD down” and “one RBD up”, were observed in our simulations, same as the observation of the virions by cryo-EM (17, 18, 21). We calculated the distance (*h*) between residues of the receptor binding domain (RBD) tip (residues 470-490 of SARS-CoV-2 and residues 457-477 of SARS-CoV) and residues at central helix (residues 986-996 of SARS-CoV-2 and residues 968-978 of SARS-CoV) (Fig. 4C). The populations of *h* in SARS-CoV systems is between 1.2 nm and 5.0 nm with a peak at 3.0 nm, while in SARS-CoV-2 systems between 1.3 nm and 6.0 nm with a peak at 3.8 nm (Fig. 4D). The distributions clearly reveal that SARS-CoV-2 has more RBD in the up state than SARS-CoV. Only Ss with RBD in the up state can bind to ACE2 and infect cells (16). So higher intrinsic flexibility may be another factor that makes SARS-CoV-2 more infectious than SARS-CoV.

**Figure 4.**
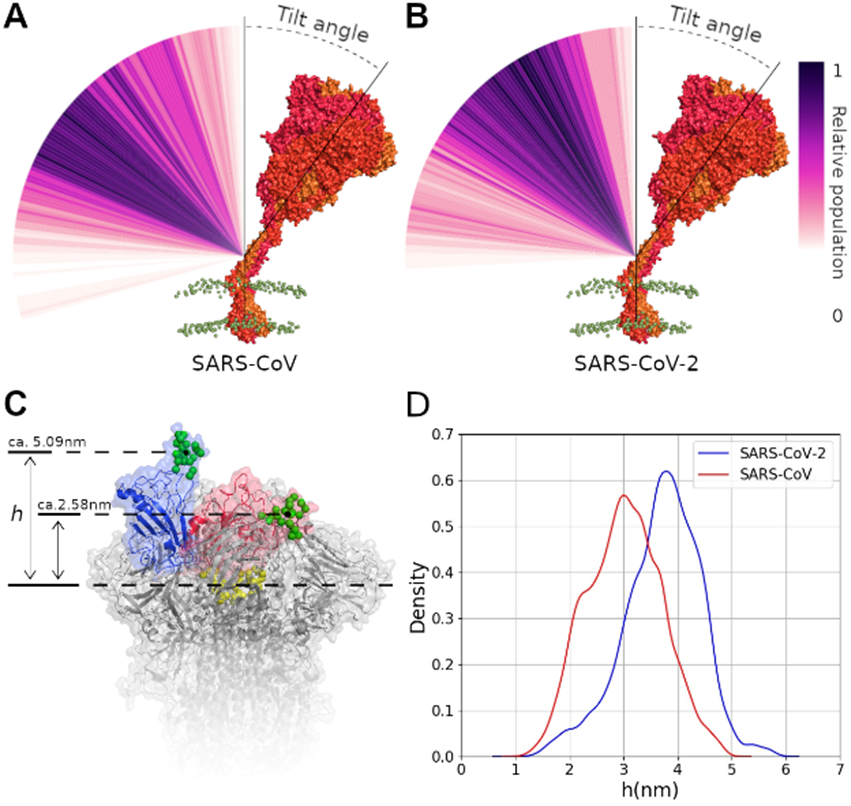
The flexibility of Ss. The distribution of the S tilt angle of SARS-CoV (A) and SARS-CoV-2 (B). (C) The distance (*h*) between the RBD tip (in green spheres) and the top of the central helix (in yellow spheres) of the closed (in red) and up states (in blue). (D) The *h* distributions of SARS-CoV-2 and SARS-CoV.

## Discussion

Microsecond MD simulations with Martini CG models were carried out to explore the dynamics and supramolecular organization of SARS-CoV and SARS-CoV-2 envelopes at experimental length scale. The presence of multiple copies of different membrane proteins allows us to get statistically significant information on PPIs and PLIs. The system complexity is comparable to those accessible by experiments such as cryo-EM. These CG models therefore provide a valuable complement to investigate the architectures of SARS-CoV and SARS-CoV-2 envelopes.

The virus envelopes maintain the spherical morphology in all simulations. Cryo-EM experiments observed both spherical and ellipsoidal virus (17, 21). The inconsistency may be because our simulations are limited to several microseconds, and we did not consider the crowding between viruses. The change in the envelope morphology is corresponding to the change of the bilayer curvature. The local curvature introduced by proteins are related to their structural symmetry (48), while the TM domains of E, M and S are small and not symmetric. The oligomerization of E in the virus membrane is still uncertain. Monomer E was adopt in our models, but some works suggested E may form homopentamer (6), which may induce local membrane curvature. However, the content of E is the least, so the influence on membrane curvature should be limited. Therefore, the ellipsoidal morphology may be mainly caused by the crowding between viruses.

In the simulations, we are able to observe the forming of S clusters and supramolecular islands by the transmembrane domains of different proteins and lipids in the envelope membrane. The interaction patterns and flexibility of S stalks show agreements with the experimental observations (17, 21, 40) and all-atom simulations (18, 49), indicating the reliability of the structural and dynamic details obtained from our simulations. The forming of S clusters can restrict the orientation of S heads, while the supramolecular islands slow down the S diffusion in the membrane. Both of these points favor the virus infection by stabilizing interactions between S and ACE2. Previous experiments have confirmed that nanocluster is required for efficient pathogen binding and internalization (50).

Our models, of course, are just approximate to the virus envelopes. In particular, the lipidomics and stoichiometric composition of structural proteins of the virus envelopes have not been accurately determined. The numbers of Ss and Ms per virion, reported in the experimental results, were speculated by assuming uniform distribution over the virus surface, and also depend on experimental conditions (41). Our current models described the virus envelopes under only one stoichiometric composition. Comparing with the stoichiometry adopted by Alvin Yu and his collaborators in their SARS-CoV-2 model (35), our models have more Ss and less Ms, which may influence the sizes of protein clusters. However, the size of the string-like protein island does not affect the mode of protein interactions.

The lipid composition of the ER-related membrane features less cholesterols and anionic lipids than the plasma membrane (PM)-related membrane (39, 51). The enrichment of cholesterols was only observed around Es, mainly due to non-specific interactions. The anionic lipids, PS and PI, make up about 15% of the inner leaflet, and are enriched around all E, M and S proteins as most proteins do (37). Therefore, the content of cholesterol and anionic lipids appears to have little effect on the PPIs in the SARS-CoV and SARS-CoV-2 envelopes.

The S protein has a dense coating of glycans, to evade the host immune system. Atomistic simulations also demonstrated that the glycans play an essential role in modulating the conformational transitions of the S protein (18, 33, 34, 49, 52). Our models in this study did not include glycans. Apparently, glycans are not involved in the PPIs of the transmembrane domains, which form the heterogeneous protein islands in the virus membrane, while the glycosylation may affect the PPIs between S heads and the size of S clusters. However, the PPI patterns in our models are consistent with these observed in the cryo-EM, as well as these from all-atom simulations, which contain four copies of glycosylated S proteins. It indicates that glycosylation may have little effect on the interaction modes between S proteins. Anyway, to get a deeper understanding of the virion structures, further studies are underway: careful parametrization of the glycosylation, and modeling and simulation of virus envelopes with glycosylation, different protein and lipid stoichiometric compositions, and expanded size dimensions.

Another possible limitation of our simulations is the use of Martini CG force field 2.2, which may limit the protein conformational changes due to the elastic network (53), such as the RBD opening in our simulations, and may tend to excessive protein aggregation because of excessive inter-protein interactions. Future simulations will try the latest Martini force fields 3.0 (54).

## Conclusions

Our Martini CG models illustrate SARS-CoV and SARS-CoV-2 envelopes at the atomistic level and the experimental complexity and scale. The structural proteins are not uniformly distributed over the envelopes. Most of Ss form oligomers with extensive interactions between their huge heads, while the transmembrane domains of structural proteins clusters into heterogeneous string-like islands, mediated by negatively charged lipids. Our simulations also revealed that the critical difference between SARS-CoV and SARS-CoV-2 envelopes lies in the S-S interaction patterns and the intrinsic flexibilities of Ss. SARS-CoV-2 Ss have more inclination to interact with each other and higher intrinsic flexibility to recognize and bind the receptors than SARS-CoV Ss, which may be two of the reasons that SARS-CoV-2 caused more infections than SARS-CoV. The structural and dynamic details of our models provide an improved understanding of the virus envelopes and could be used for further studies, such as drug design, and fusion and fission processes.

## Methods

### Simulation system setup

A coarse-grained (CG) vesicle with an inner diameter of approximate 45 nm, that is an outer diameter of about 53 nm, was modeled by CHARMM-GUI Martini Maker (55). According to the coronavirus biogenesis, the virus envelope buds from the endoplasmic reticulum Golgi intermediate compartment (ERGIC) (56), whose membrane shares properties with the ER-NE-*cis*-Golgi lipid territory. Therefore, the lipid composition of the virus envelope is based on the lipid composition of membranes of the ER-NE-*cis*-Golgi lipid territory (39) (Fig. 1 and Table S1). The vesicle was equilibrated by a 2 μs MD simulation with its radius of gyration stabilizing at ∼22.3 nm.

The genes of the E and M proteins are conserved in SARS-CoV and SARS-CoV-2 (Fig. S1), so the same structures of M and E were used in both models. The oligomerization of E differs from monomer (57, 58) to pentamer (6, 58) under different experimental conditions without M and S proteins, but is still not demonstrated in the CoV membrane. We chose to involve monomers of E in our models. The monomer structure of E was extracted from the SARS-CoV E protein pentameric structure (PDB:5×29) (58), while the M structure downloaded from the I-TASSER website (22) was used with the adjustment of the carboxy-terminal domain (CTD) to the virus lumen (Fig. S3). The adjusted conformation is consistent with the newly released conformation on the I-TASSER website.

The main difference between SARS-CoV and SARS-CoV-2 in gene sequence is centered on the S protein. We used the closed state prefusion structure of SARS-CoV S (PDB:5×58) (59) and SARS-CoV-2 S (PDB:6VXX) (60) respectively. For the SARS-CoV S, the missing TM region of S was taken from the predicted structure from I-TASSER website (22) and other missing fragments such as HR2 from other structures (PDBs: 6NB6, 6B3O, 2FXP). And the missing fragments of SARS-CoV-2 S was modeled based on this complete SARS-CoV S structure. The protonation states of all residues and lipids uses their states at pH=7. There is no clear consensus on the stoichiometric composition of structural proteins of the envelopes so far. Here, a total of 90 proteins (15 Es, 30 Ss and 45 Ms) were inserted into the vesicles randomly, and then the lipid molecules within 2 Å of proteins were removed, resulting in 16,000 lipid molecules left. Then, the envelope models were solvated in a dodecahedron-shaped box with Martini water and counter ions to neutralize the overall charge and 0.15M NaCl, resulting in simulation systems of about 7.3 million Martini beads.

### Molecular Dynamics Simulations

All CG MD simulations were performed using GROMACS version 2019.2 (61) and with Martini 2.2 force field parameters (62, 63) and the ELNEDYN elastic network model on monomers, using a force constant of 500 kJ/(mol•nm^2^) and a distance cutoff of 0.9 nm. The temperature was maintained at 310 K using a velocity-rescaling thermostat (64) with a time constant for coupling of 1 ps. Proteins and lipids, and solvent were coupled separately to the temperature bath. An isotropic pressure of 1 bar was maintained with the (65) barostat, with a compressibility of 4.5×10^−5^ bar^-1^, and a relaxation time constant of 5 ps. A cutoff of 12 Å was used for van der Waals and electrostatic interactions with a switching function from 10 Å for van der Waals. The systems were minimized for 15000 steps with the steepest descent method, and then equilibrated by short 3.6 ns NVT simulations with incremental time steps of 2fs, 5fs 10fs, 15fs and 20fs. Finally, the production simulations were performed with the NPT ensemble and a time step of 20 fs. Both simulation systems were carried out for multiple replicas, for a total of 60 μs (Table S2). All the analyses were performed using VMD (66), which is also used for visualization and figure rendering.

## ASSOCIATED CONTENT

### Supporting Information

The Supporting Information is available free of charge on the website at DOI:

Supplementary methods, tables and figures (PDF)

Movie S1. One of the SARS-CoV-2 simulations. The color scheme is the same as Figure 1.

### Notes

The authors declare no competing financial interest.

## Acknowledgments

This work was supported by the Natural Sciences and Engineering Research Council (Canada), the Canada Research Chairs program, the National Natural Science Foundation of China (No. 31971176 and 31800616) and the Fundamental Research Funds for the Central Universities (No. A03018023601045). Calculations were run on Compute Canada facilities, funded by the Canada Foundation for Innovation and partners.

