## Supporting information for "The supramolecular organization of SARS-CoV and SARS-CoV-2 virions revealed by coarse-grained models of intact virus envelopes"

### Methods

#### Trajectory analysis

**Lipid depletion-enrichment index.** For a generic lipid type L, defining the ration of lipid within a given cut-off $x$ (termed Ratio(L)$_x$), the ratio of the lipid L with respect to bulk (termed Ratio(L)$_{bulk}$) is calculated as following:

$$Ratio(L)_x = \frac{(no.L)_x}{(tot.no.lipids)_x}$$

$$Ratio(L)_{bulk} = \frac{tot.no.(L)}{(tot.no.lipids)},$$

where no.L is the number of the lipid L and tot.no.lipids is number of the total lipids. The depletion-enrichment (D-E) index of the lipid L is their ratio. In our analyses, lipid headgroups were used to calculate the D-E index within a cut-off of 0.7nm from any bead of proteins. The ROH bead was chosen for cholesterol, while GL1 or AM1 beads were used for all the other lipid types. The two leaflets were combined together.

Table S1. Lipid composition of viral envelopes.

| Outer leaflet | | | Inner leaflet | | |
|---|---|---|---|---|---|
| Lipid type | No. | Percentage | Lipid type | No. | Percentage |
| CHOL | 420 | 4.63% | CHOL | 376 | 5.20% |
| DAPC | 62 | 0.68% | DAPC | 28 | 0.39% |
| DOPC | 178 | 1.96% | DOPC | 89 | 1.23% |
| PAPC | 490 | 5.40% | PAPC | 230 | 3.18% |
| PEPC | 126 | 1.39% | PEPC | 63 | 0.87% |
| PIPC | 3097 | 34.15% | PIPC | 1471 | 20.36% |
| POPC | 2105 | 23.21% | POPC | 1007 | 13.94% |
| PUPC | 122 | 1.35% | PUPC | 51 | 0.71% |
| DAPE | 81 | 0.89% | DAPE | 373 | 5.16% |
| DOPE | 51 | 0.56% | DOPE | 217 | 3.00% |
| DUPE | 24 | 0.26% | DUPE | 106 | 1.47% |
| PAPE | 142 | 1.57% | PAPE | 587 | 8.12% |
| PIPE | 103 | 1.14% | PIPE | 428 | 5.92% |
| POPE | 150 | 1.65% | POPE | 638 | 8.83% |
| PQPE | 26 | 0.29% | PQPE | 112 | 1.55% |
| PUPE | 53 | 0.58% | PUPE | 221 | 3.06% |
| BNSM | 36 | 0.40% | BNSM | 17 | 0.24% |
| DBSM | 25 | 0.28% | DBSM | 10 | 0.14% |
| DPSM | 111 | 1.22% | DPSM | 52 | 0.72% |
| DXSM | 47 | 0.52% | DXSM | 22 | 0.30% |
| PGSM | 7 | 0.08% | PGSM | 4 | 0.06% |
| PNSM | 70 | 0.77% | PNSM | 36 | 0.50% |
| POSM | 7 | 0.08% | POSM | 2 | 0.03% |
| XNSM | 50 | 0.55% | XNSM | 19 | 0.26% |
| APC | 206 | 2.27% | DAPS | 8 | 0.11% |
| IPC | 211 | 2.33% | DUPS | 8 | 0.11% |
| OPC | 239 | 2.64% | PAPS | 180 | 2.49% |
| PPC | 750 | 8.27% | PIPS | 31 | 0.43% |
| UPC | 81 | 0.89% | POPS | 81 | 1.12% |
| | | | PQPS | 18 | 0.25% |
| | | | PUPS | 71 | 0.98% |
| | | | PAPI | 181 | 2.50% |
| | | | PIPI | 195 | 2.70% |
| | | | POPI | 215 | 2.98% |
| | | | PUPI | 79 | 1.09% |

Table S2. Simulation information.

| Simulation System | | System Size | Simulation Length |
|---|---|---|---|
| SARS-CoV | SARS-CoV-md1 | 7.27M | 12.0 μs |
| | SARS-CoV-md2 | 7.27M | 12.0 μs |
| | SARS-CoV-md3 | 7.27M | 8.0 μs |
| SARS-CoV-2 | SARS-CoV-2-md1 | 7.25M | 5.0 μs |
| | SARS-CoV-2-md2 | 7.25M | 3.0 μs |
| | SARS-CoV-2-md3 | 7.25M | 5.0 μs |
| | SARS-CoV-2-md4 | 7.25M | 6.5 μs |
| | SARS-CoV-2-md5 | 7.25M | 8.0 μs |

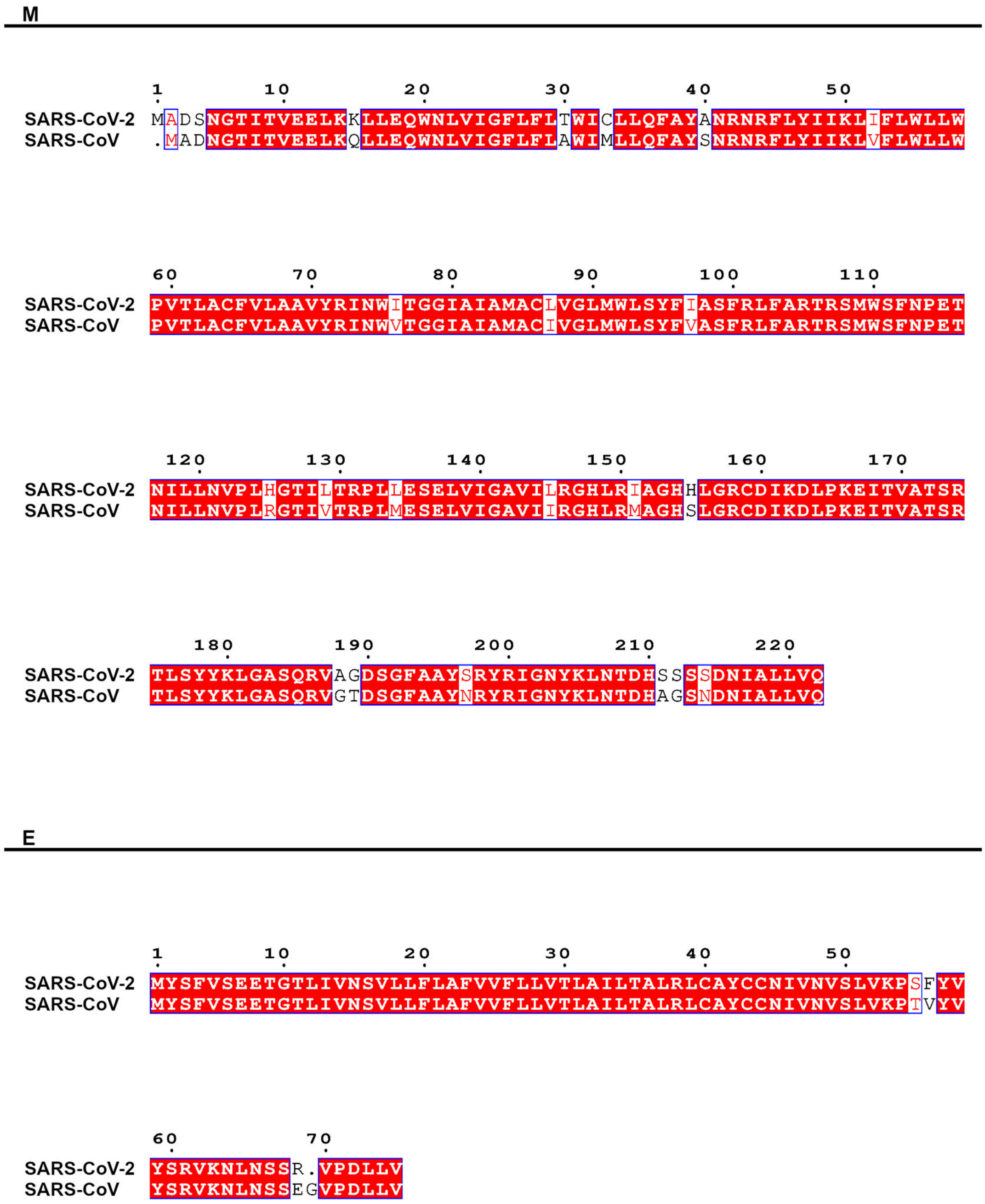


Figure S1. The sequence alignment of M and E proteins of SARS-CoV and SARS-CoV-2.

**S**

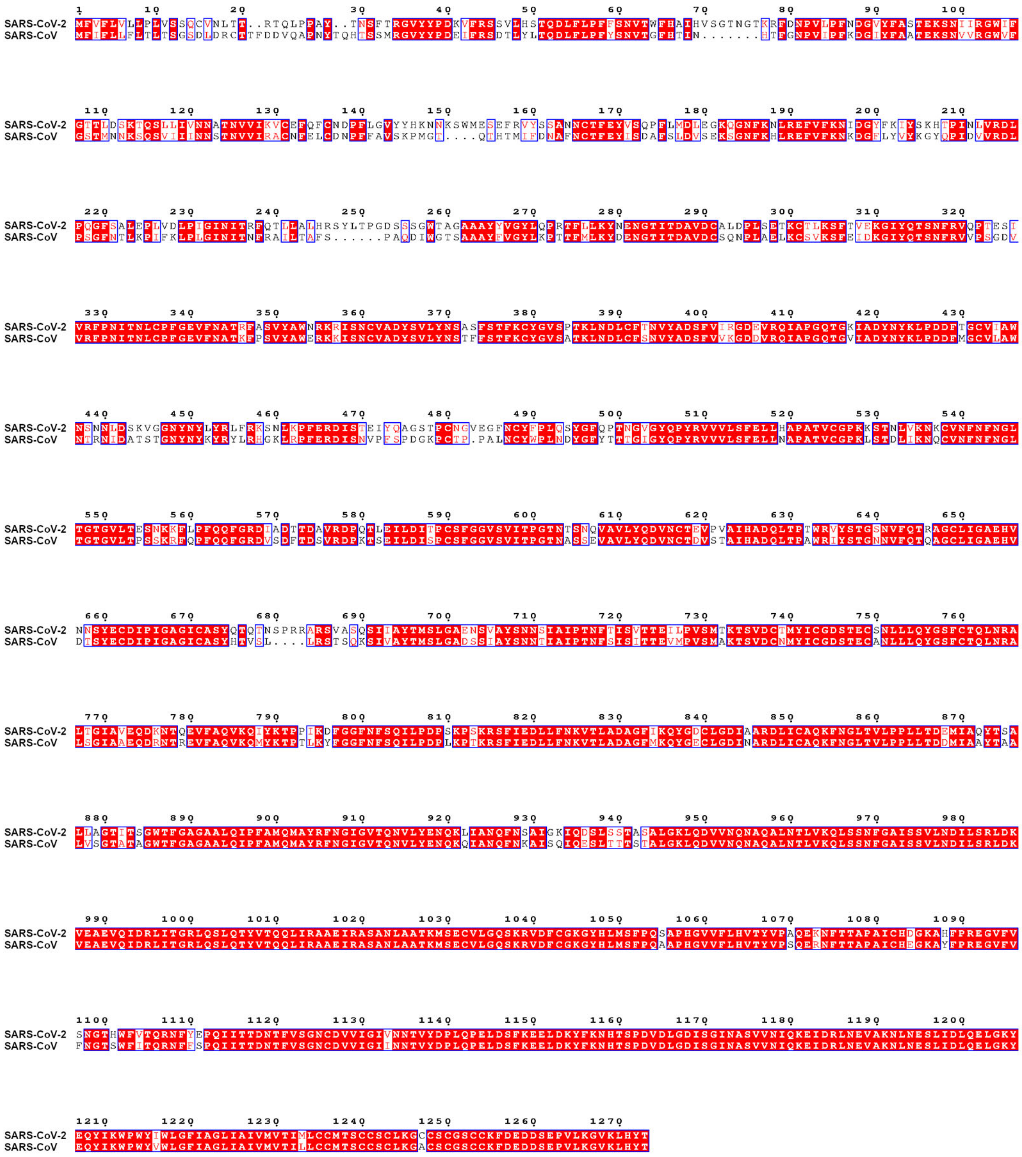


Figure S2. The sequence alignment of S proteins of SARS-CoV and SARS-CoV-2. Each monomer consists of two subunits: S1 (1-685) and S2 (686-1273). The main difference between the two sequences is concentrated on S1 subunit.

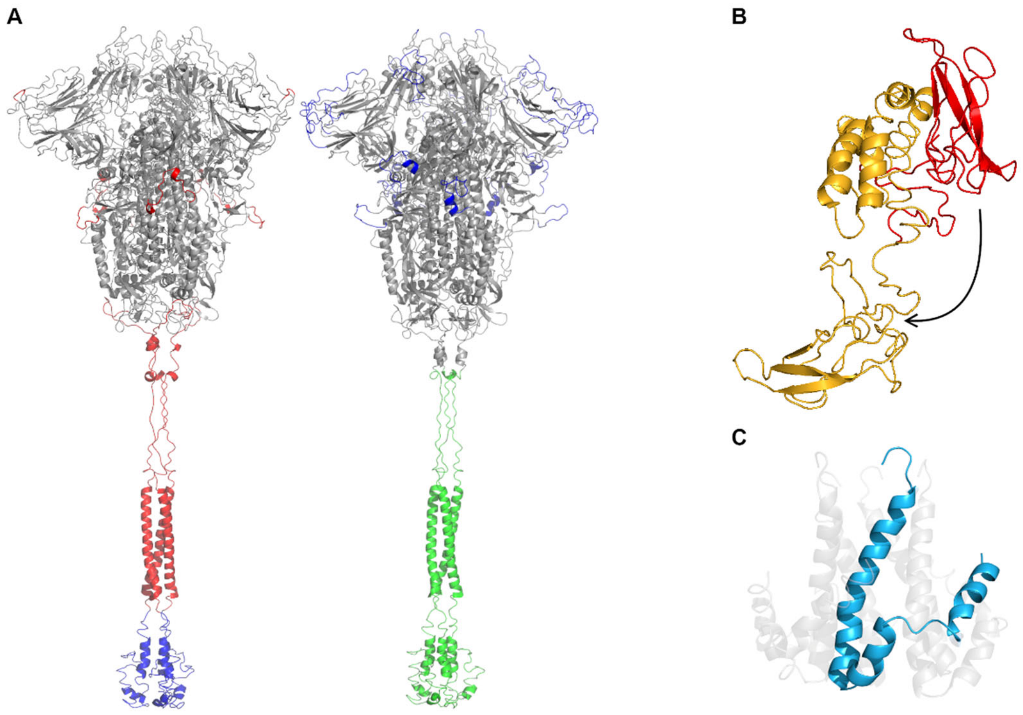


Figure S3. The initial structures of S (A), M (B) and E (C). Different S structures were used for SARS-CoV (A, left) and SARS-CoV-2 (A, right). (A) The structural model of SARS-CoV is based on the cryo-EM structure (PDB ID: 5X58), and the missing fragments are taken from other cryo-EM structures (PDB IDs: 6CRX, 6NB6, 6B3O and 2FXP, in red) and the predicted structure by I-TASSER (in blue). The structural model of SARS-CoV-2 is based on the cryo-EM structure (PDB ID: 6VXX), and the missing fragments are taken from the predicted structure by I-TASSER (in blue) and the structural model obtained by homology modeling using the SARS structure as template. (B) In the predicted M structure by I-TASSER, the endodomain, in red, was located at the position of the lipid bilayer. So, the endodomain was adjusted to the intracellular space, showing in yellow.

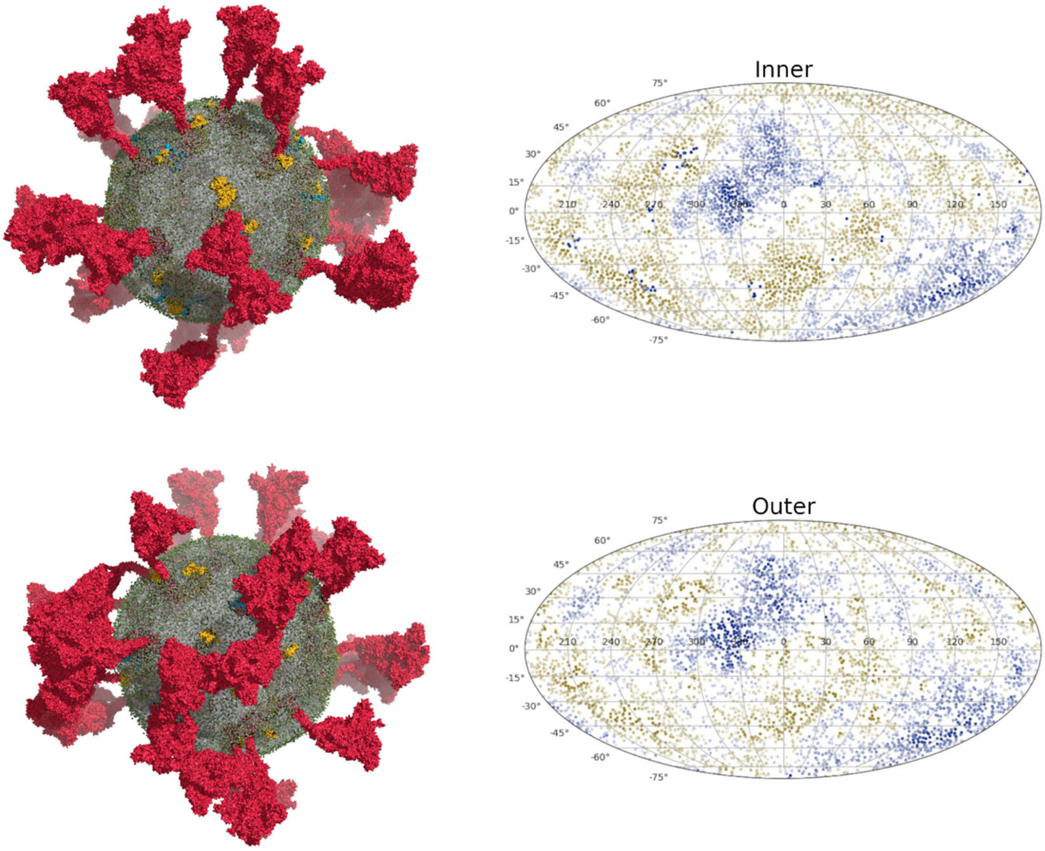


Figure S4. The side view of equilibrated CG models of SARS-CoV-2 and Mollweide projection maps of the distances between lipid heads and the vesicle’s center of mass from simulations SARS-CoV-2-md1.

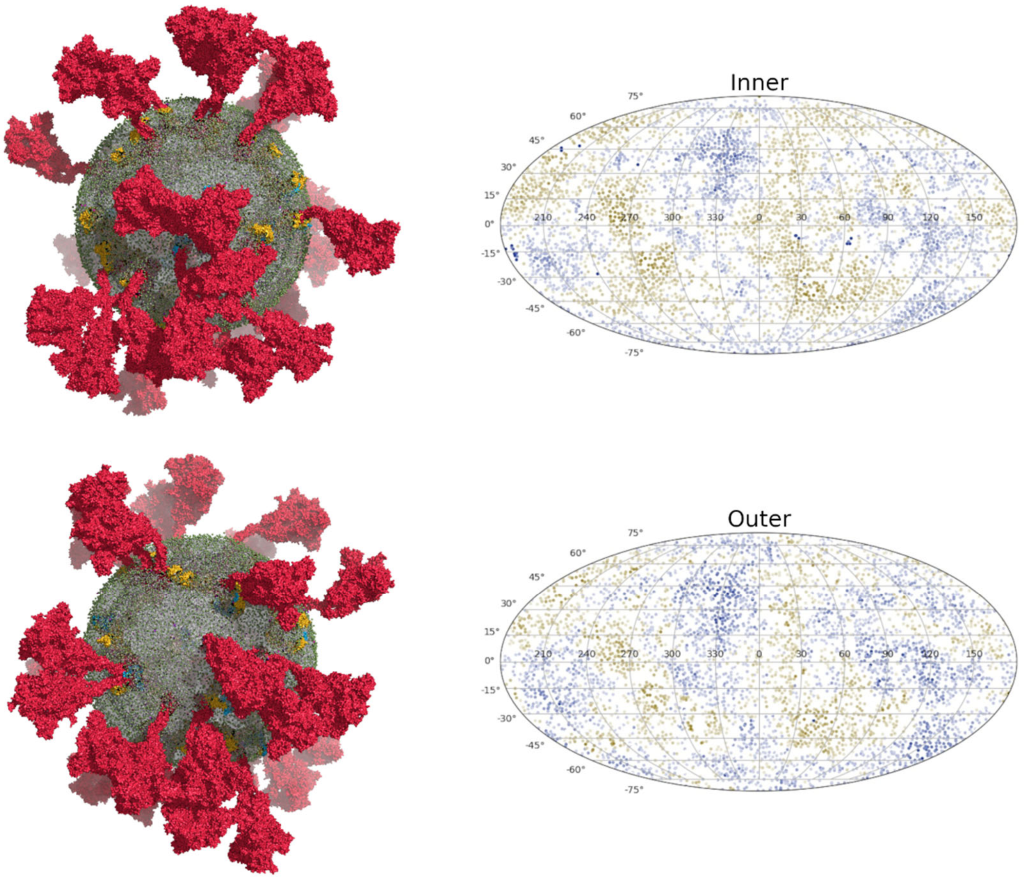


Figure S5. The side view of equilibrated CG models of SARS-CoV-2 and Mollweide projection maps of the distances between lipid heads and the vesicle’s center of mass from simulations SARS-CoV-2-md2.

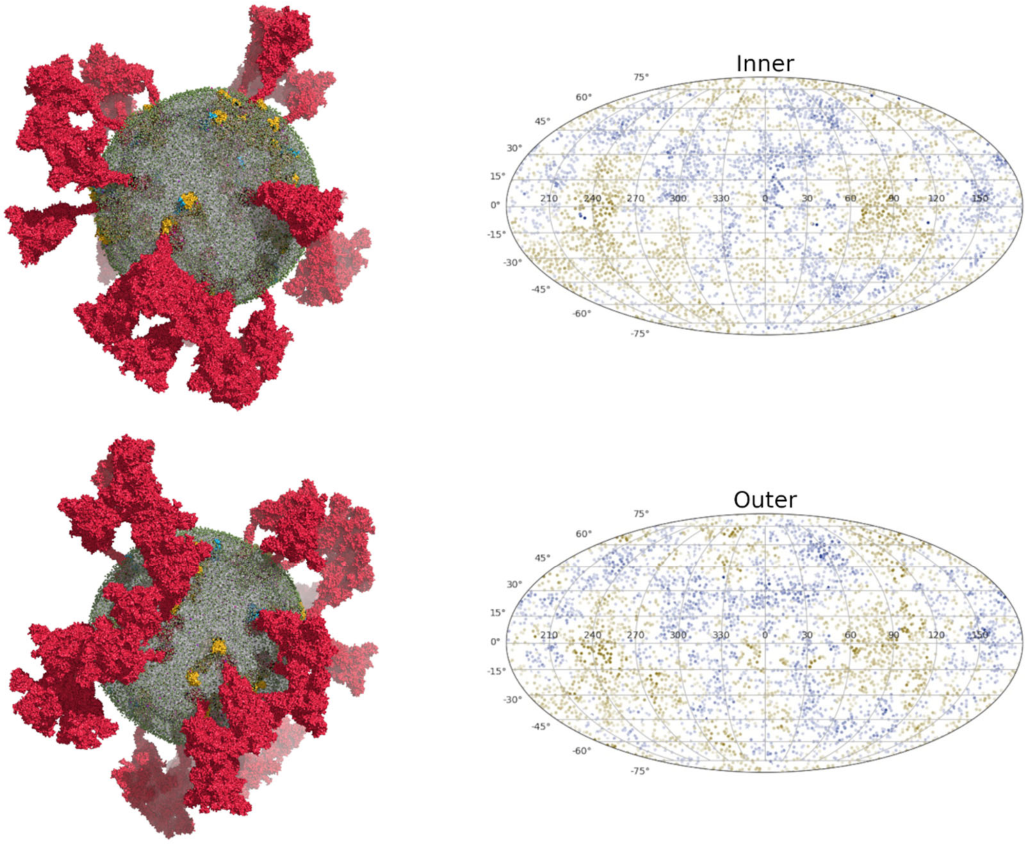


Figure S6. The side view of equilibrated CG models of SARS-CoV-2 and Mollweide projection maps of the distances between lipid heads and the vesicle’s center of mass from simulations SARS-CoV-2-md3.

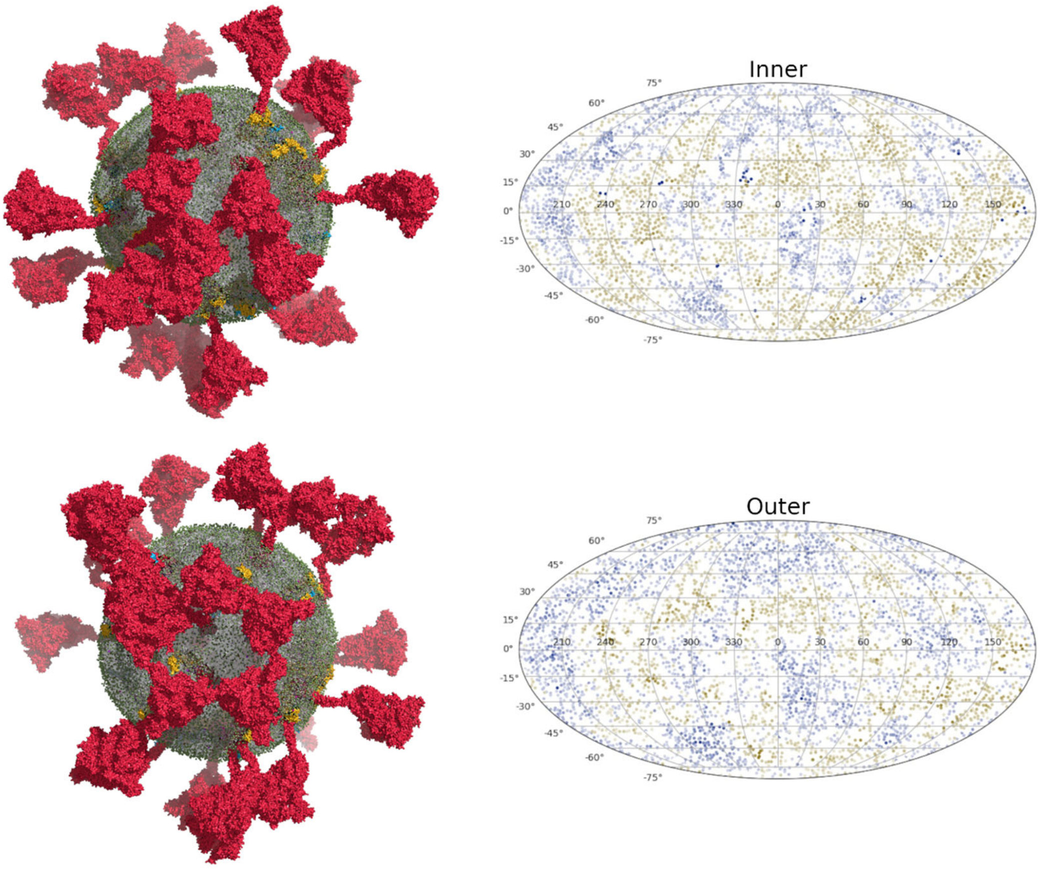


Figure S7. The side view of equilibrated CG models of SARS-CoV-2 and Mollweide projection maps of the distances between lipid heads and the vesicle’s center of mass from simulations SARS-CoV-2-md4.

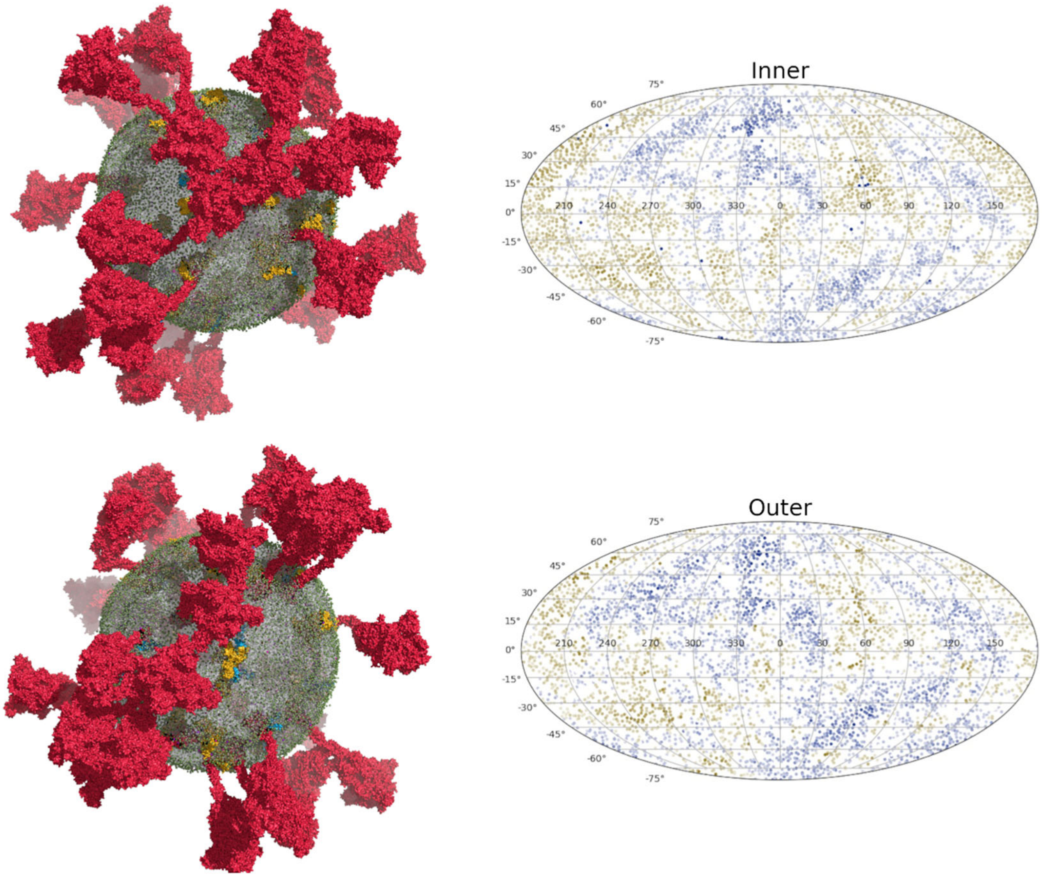


Figure S8. The side view of equilibrated CG models of SARS-CoV-2 and Mollweide projection maps of the distances between lipid heads and the vesicle's center of mass from simulations SARS-CoV-2-md5.

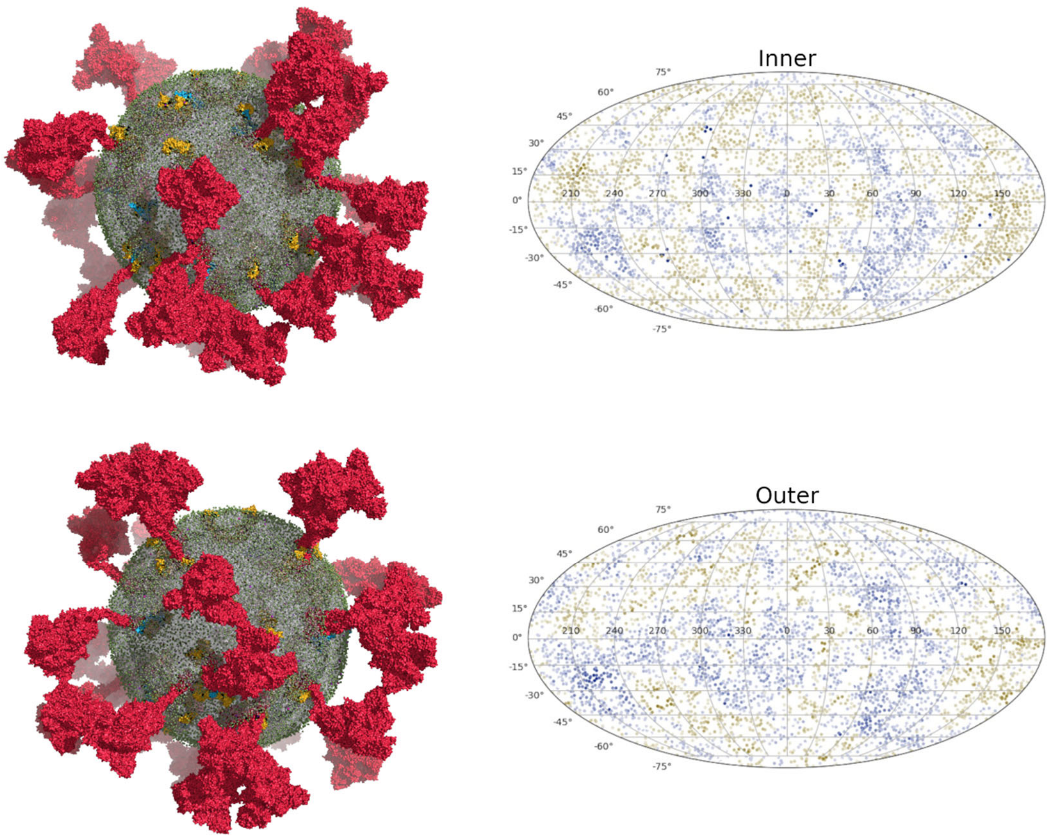


Figure S9. The side view of equilibrated CG models of SARS-CoV and Mollweide projection maps of the distances between lipid heads and the vesicle's center of mass from simulations SARS-CoV-2-md1.

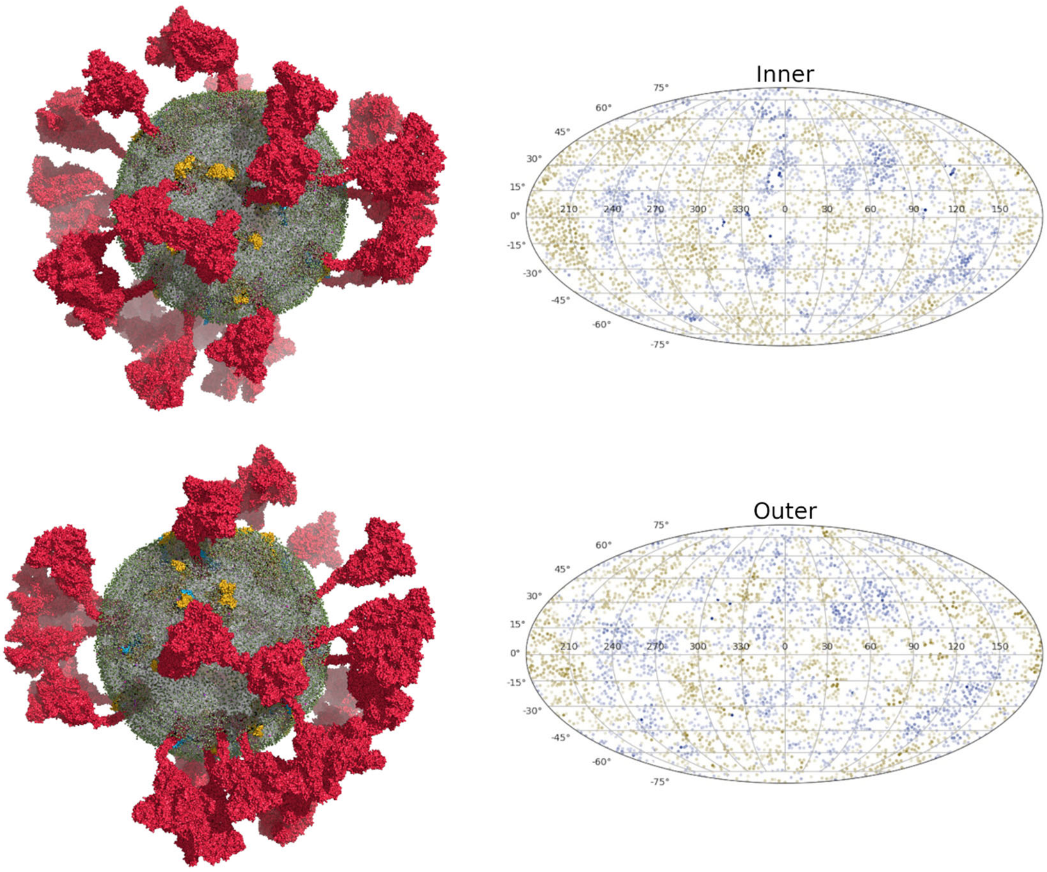


Figure S10. The side view of equilibrated CG models of SARS-CoV and Mollweide projection maps of the distances between lipid heads and the vesicle's center of mass from simulations SARS-CoV-2-md2.

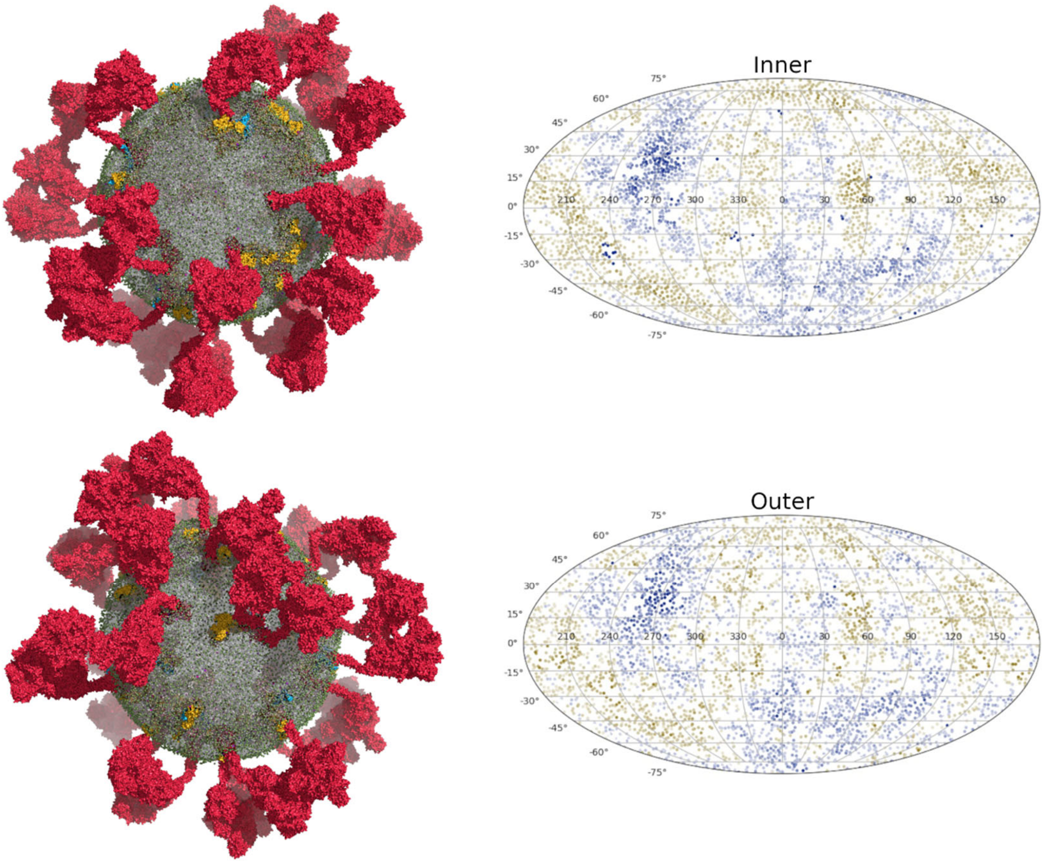


Figure S11. The side view of equilibrated CG models of SARS-CoV and Mollweide projection maps of the distances between lipid heads and the vesicle’s center of mass from simulations SARS-CoV-2-md3.

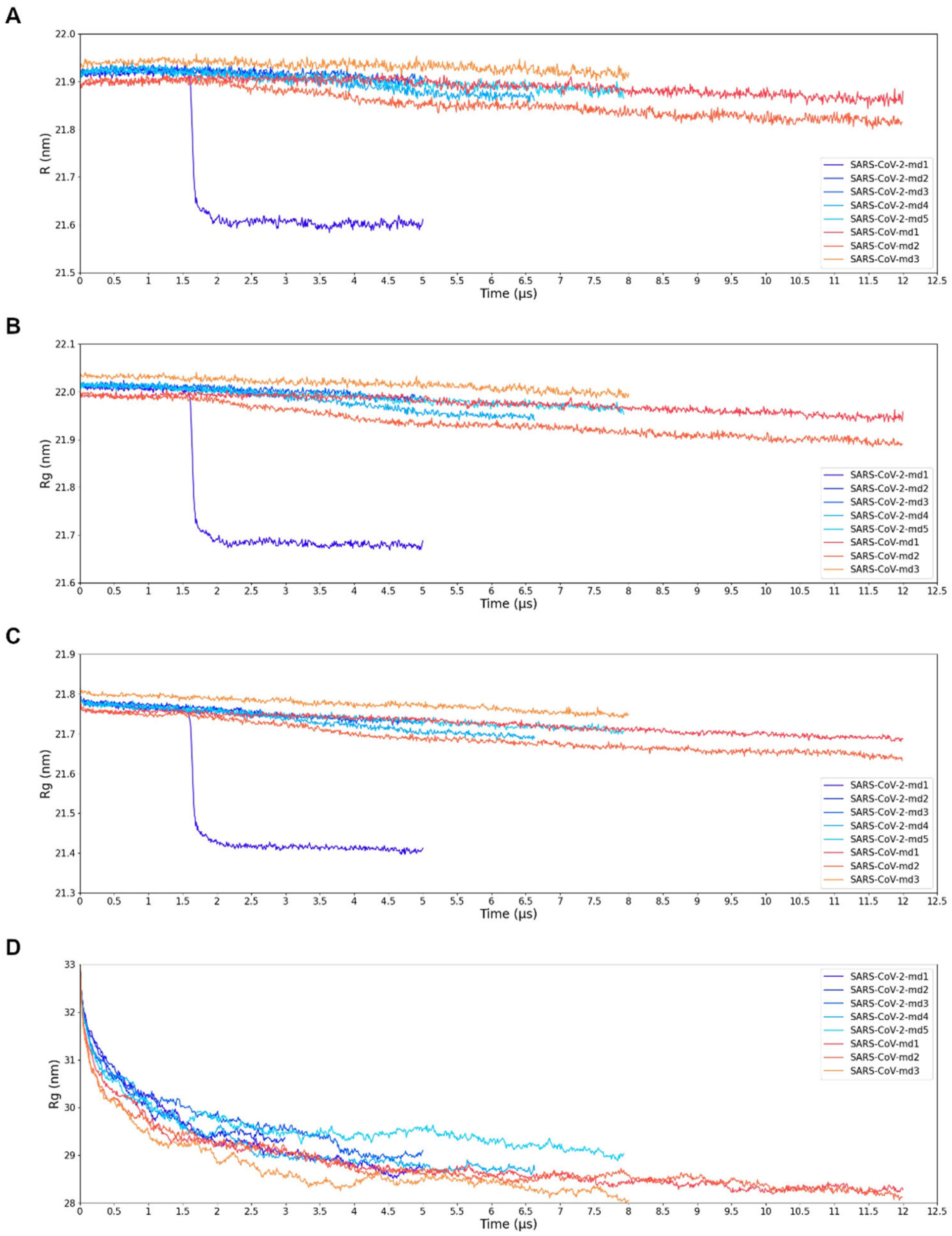


Figure S12. The variation of average curvature radius (*R*) (A), radius of gyration ($R_g$) of envelopes without proteins (B), radius of gyration ($R_g$) of envelopes without the ectodomains of S (C), and $R_g$ of intact envelopes (D) over the simulation time.

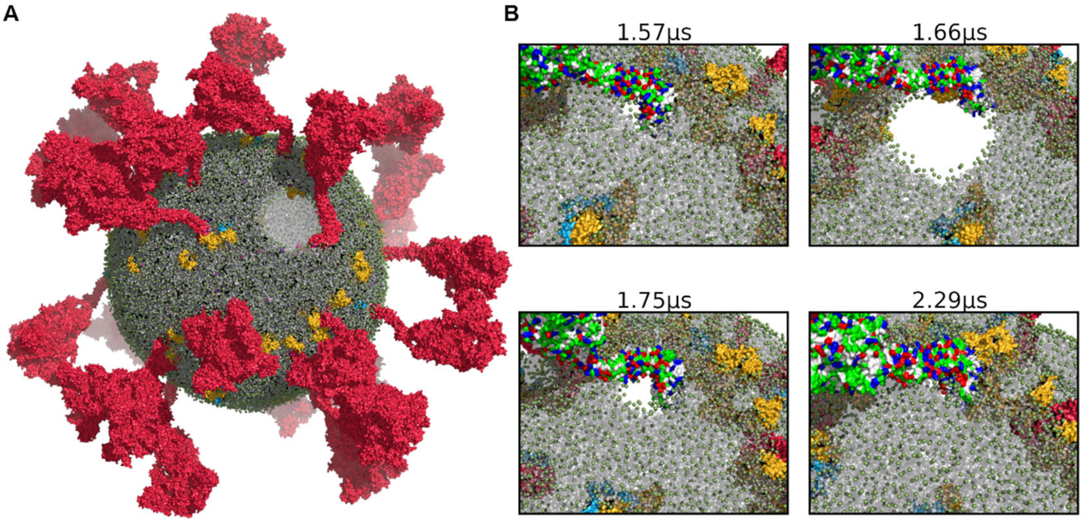


Figure S13. (A) The global view of the hole induced by one S's laying down. (B) Snapshots of the hole forming and disappearing. The S was colored according to the physicochemical properties of amino acids: polar residues in green, basic residues in blue, acidic residues in red and nonpolar residues in white.

The stalk of one S falls on the surface of the lipid bilayer, pulling the C-terminal endodomain into the lipid bilayer. The hydrophobic mismatch and electrostatic repulsion push the lipids away, causing a hole with a diameter of as large as about 9.0 nm and a decrease of the area per lipid. After rearrangement of surrounding lipid types, the hole gets smaller and eventually disappears. The formation of a hole by S's laying down leads to a more compact lipid bilayer.

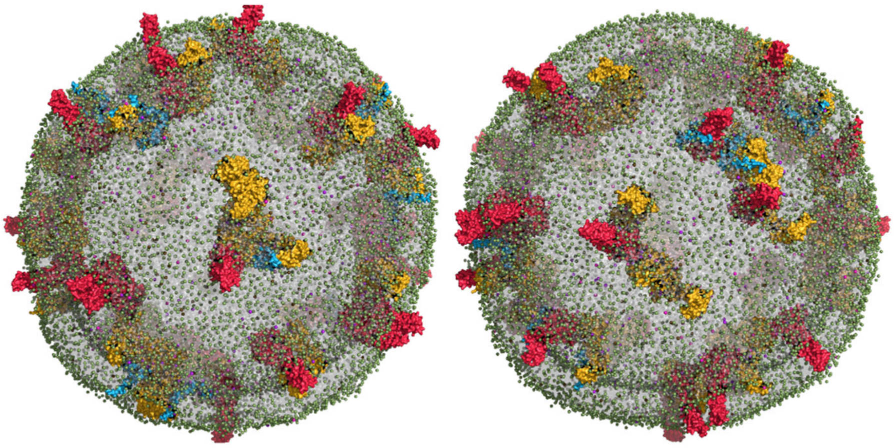

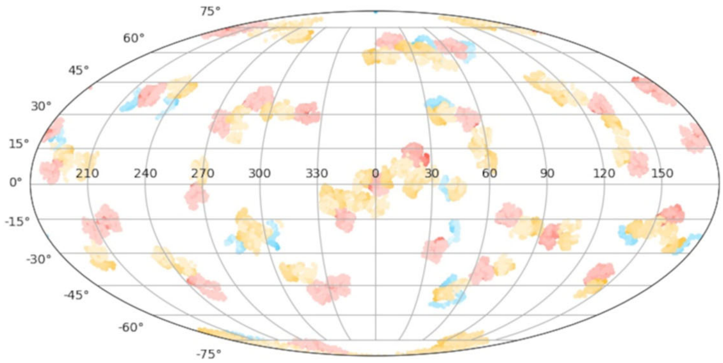


Figure S14. The equilibrated CG models of SARS-CoV-2 envelopes visualizing without the S ectodomains, and its Mollweide projection map of protein distribution from simulations SARS-CoV-2-md1.

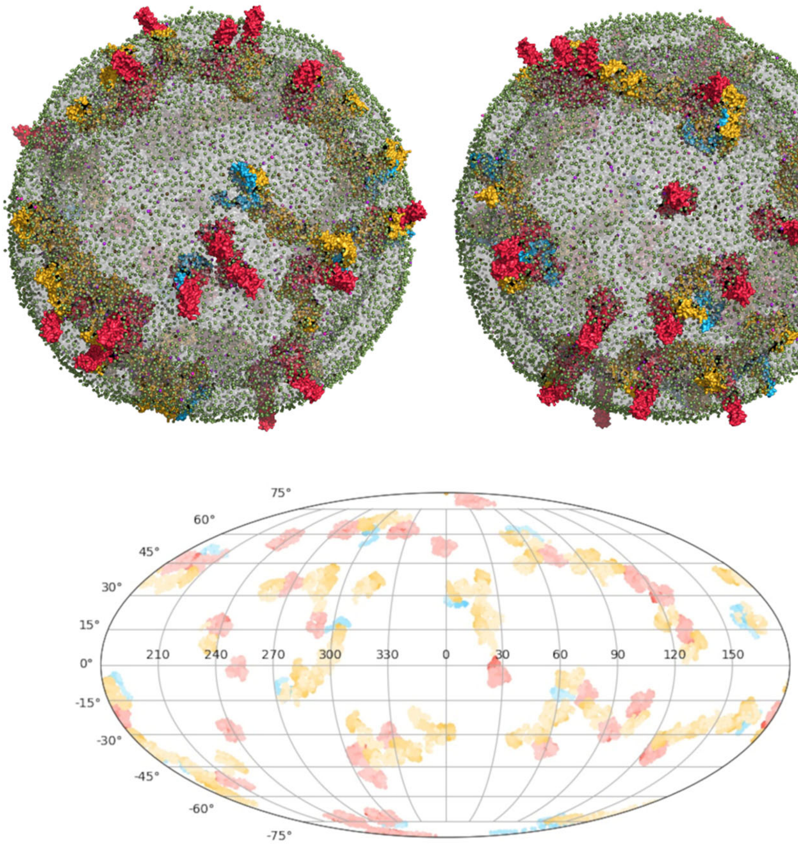


Figure S15. The equilibrated CG models of SARS-CoV-2 envelopes visualizing without the S ectodomains, and its Mollweide projection map of protein distribution from simulations SARS-CoV-2-md2.

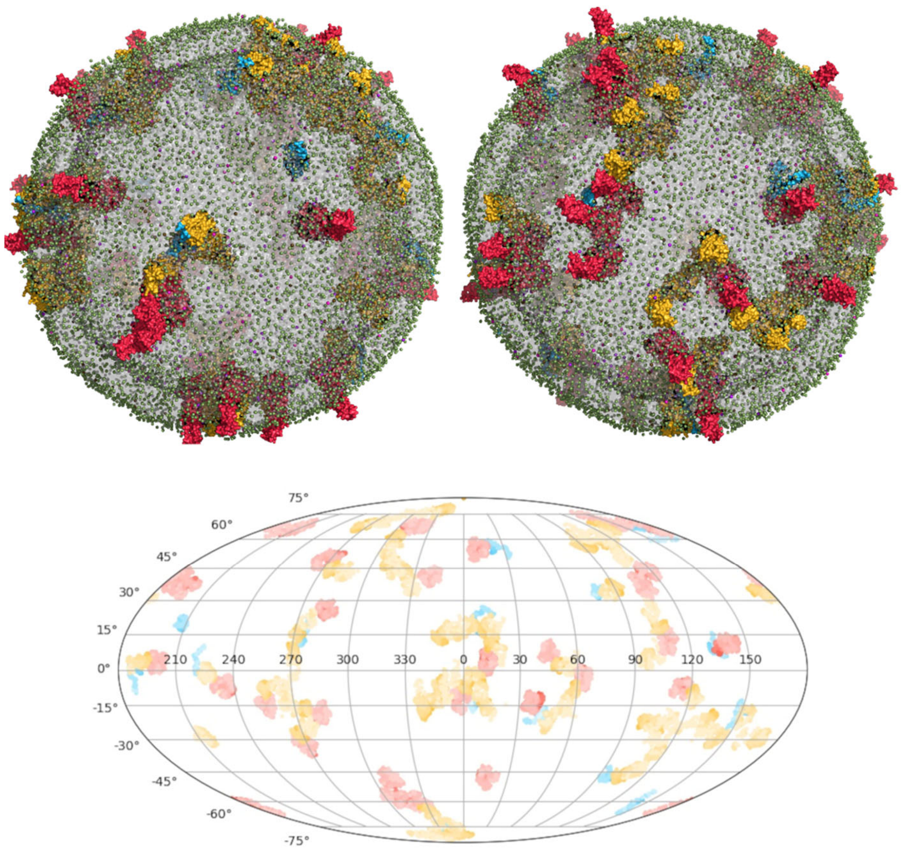


Figure S16. The equilibrated CG models of SARS-CoV-2 envelopes visualizing without the S ectodomains, and its Mollweide projection map of protein distribution from simulations SARS-CoV-2-md3.

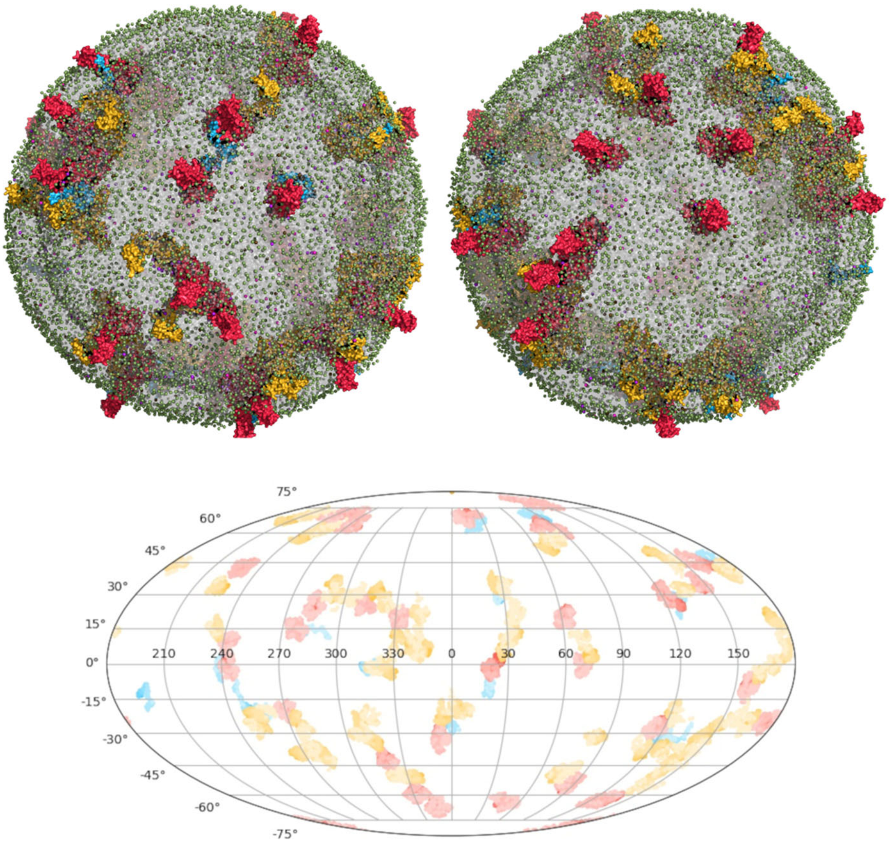


Figure S17. The equilibrated CG models of SARS-CoV-2 envelopes visualizing without the S ectodomains, and its Mollweide projection map of protein distribution from simulations SARS-CoV-2-md4.

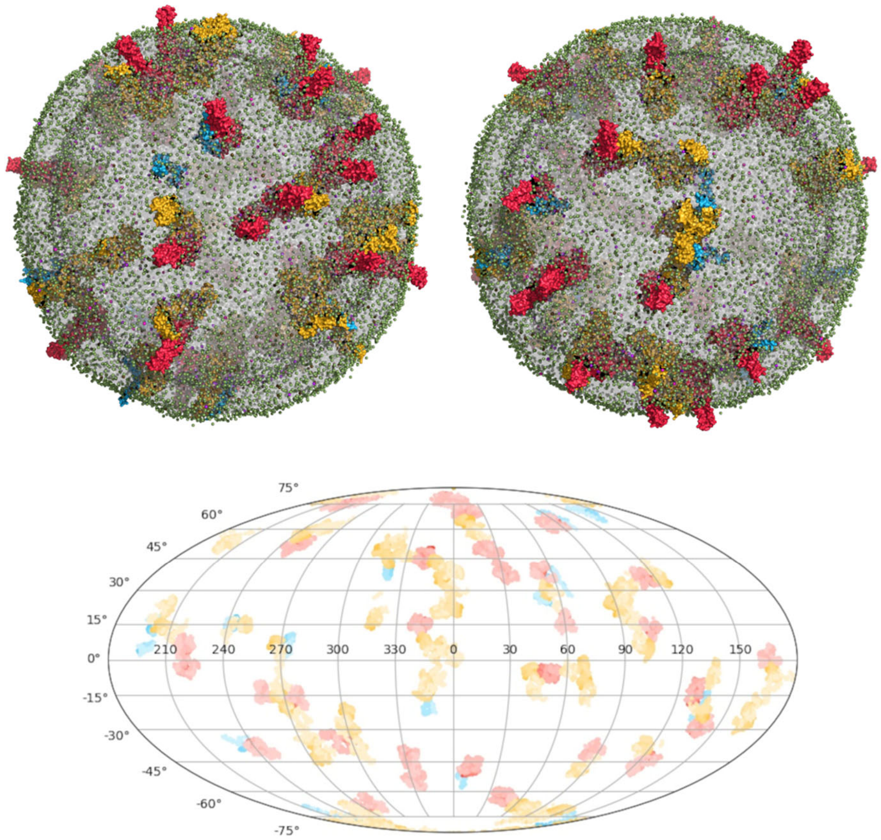


Figure S18. The equilibrated CG models of SARS-CoV-2 envelopes visualizing without the S ectodomains, and its Mollweide projection map of protein distribution from simulations SARS-CoV-2-md5.

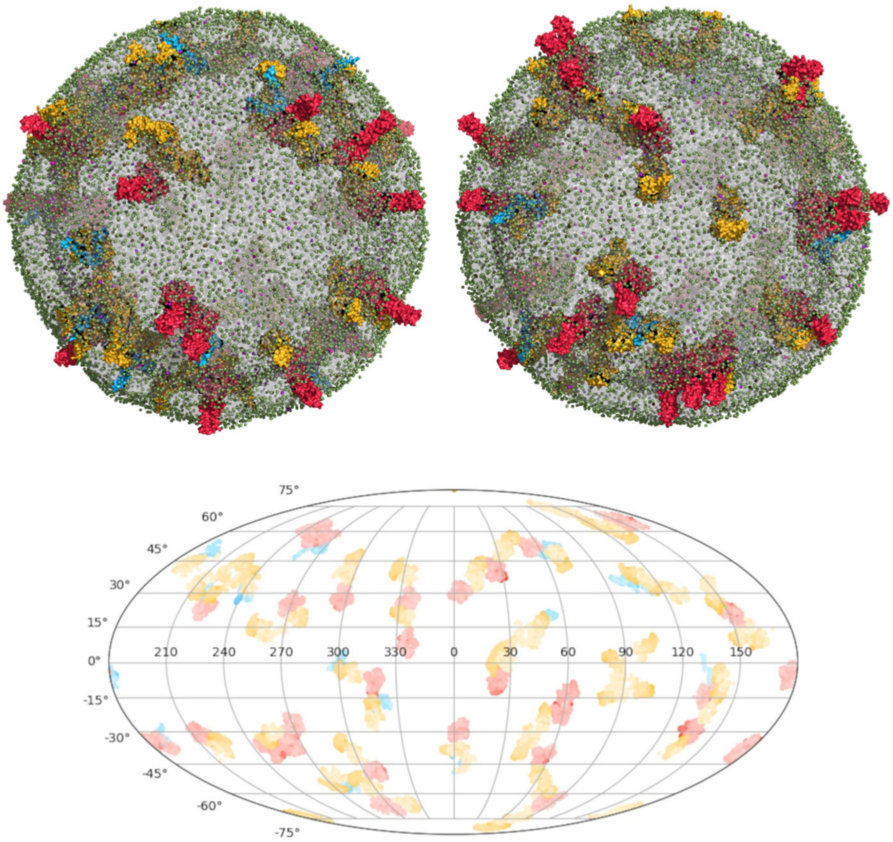


Figure S19. The equilibrated CG models of SARS-CoV envelopes visualizing without the S ectodomains, and its Mollweide projection map of protein distribution from simulations SARS-CoV-md1.

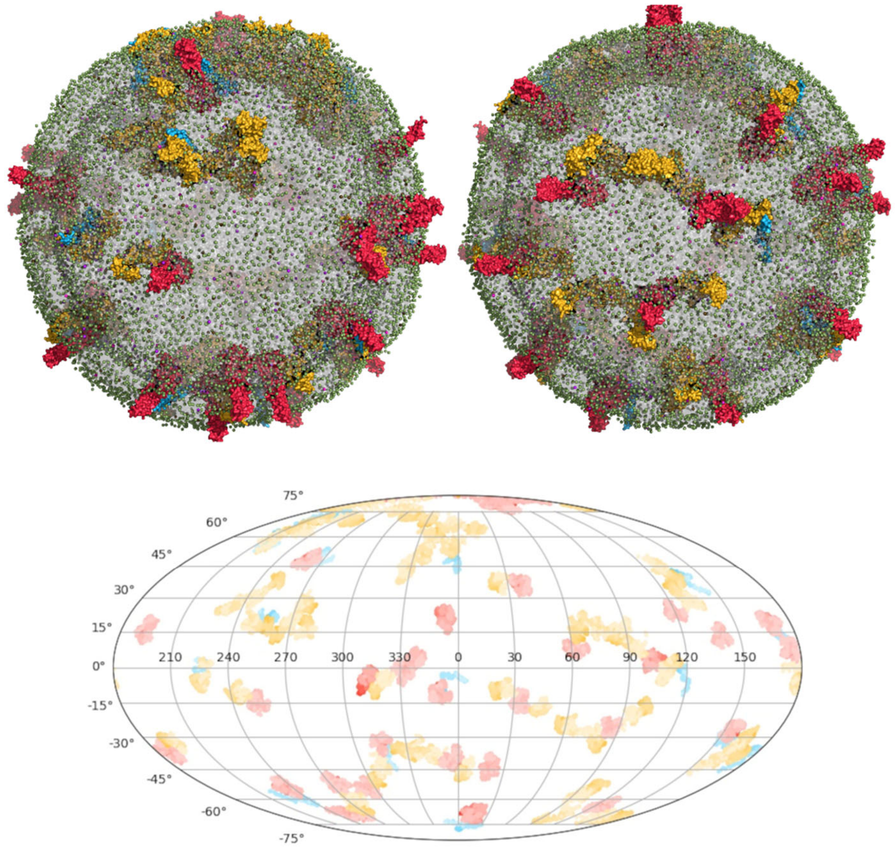


Figure S20. The equilibrated CG models of SARS-CoV envelopes visualizing without the S ectodomains, and its Mollweide projection map of protein distribution from simulations SARS-CoV-md2.

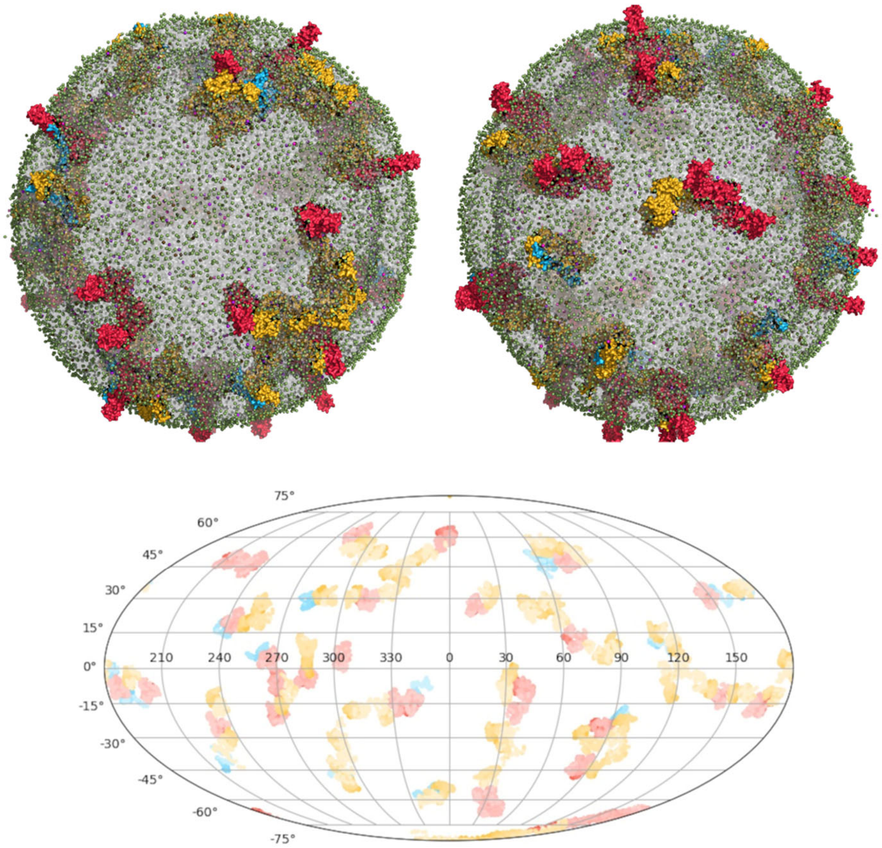


Figure S21. The equilibrated CG models of SARS-CoV envelopes visualizing without the S ectodomains, and its Mollweide projection map of protein distribution from simulations SARS-CoV-md3.

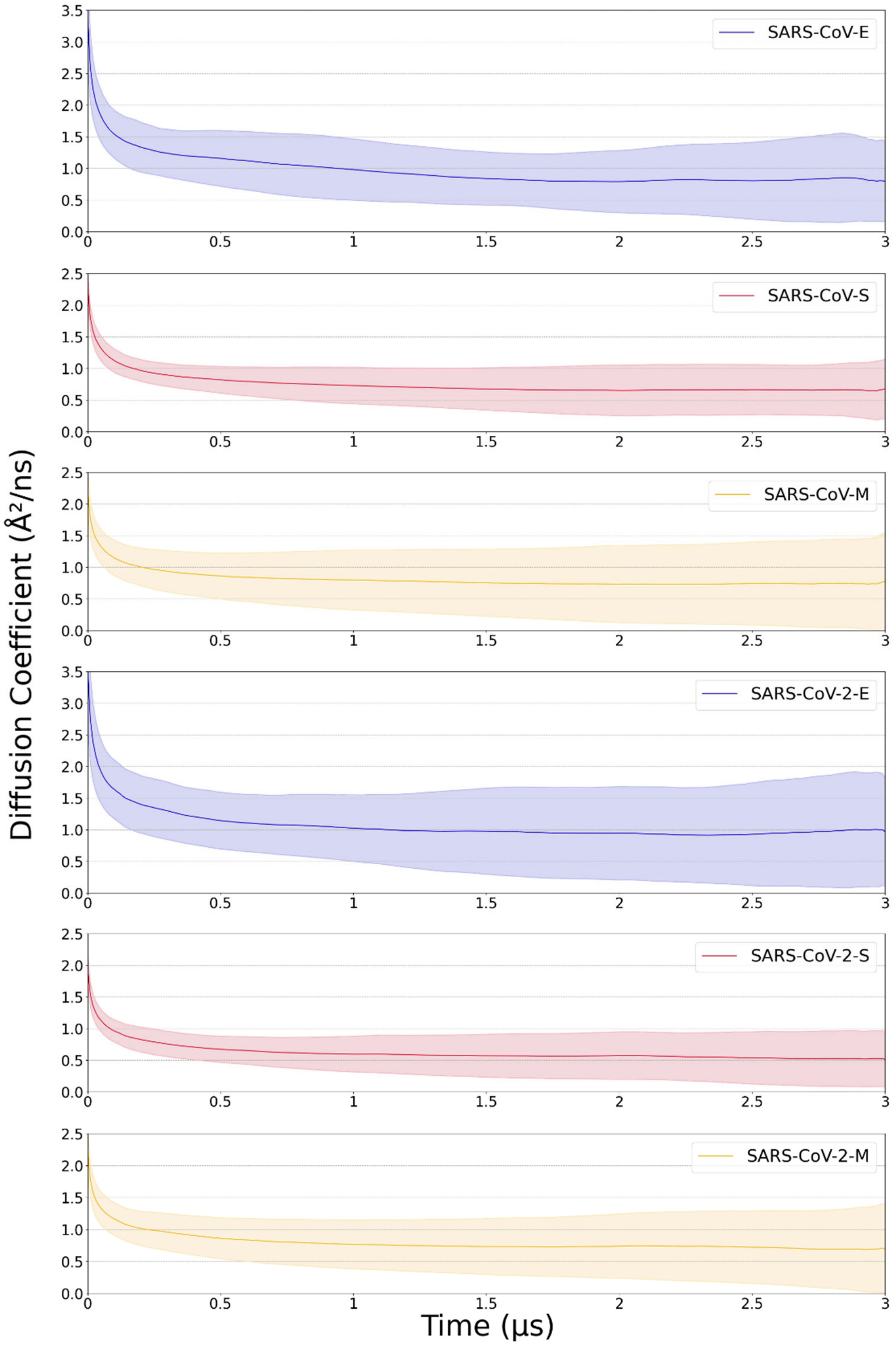


Figure S22. Lateral diffusion coefficient of structural proteins, obtained from fitting to the mean-square displacement.

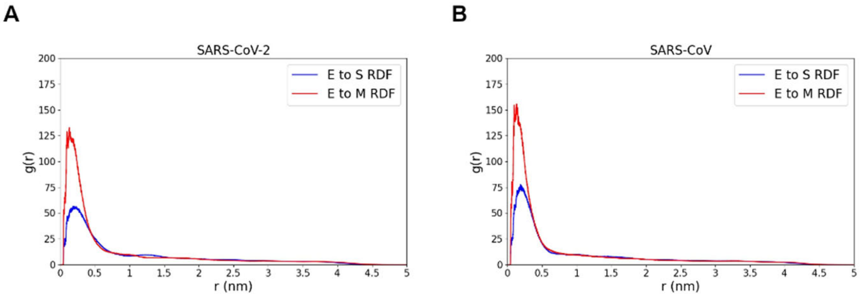


Figure S23. The radial distribution function (g(r)) of E and S, and E and M for SARS-CoV-2 (A) and SARS-CoV (B).

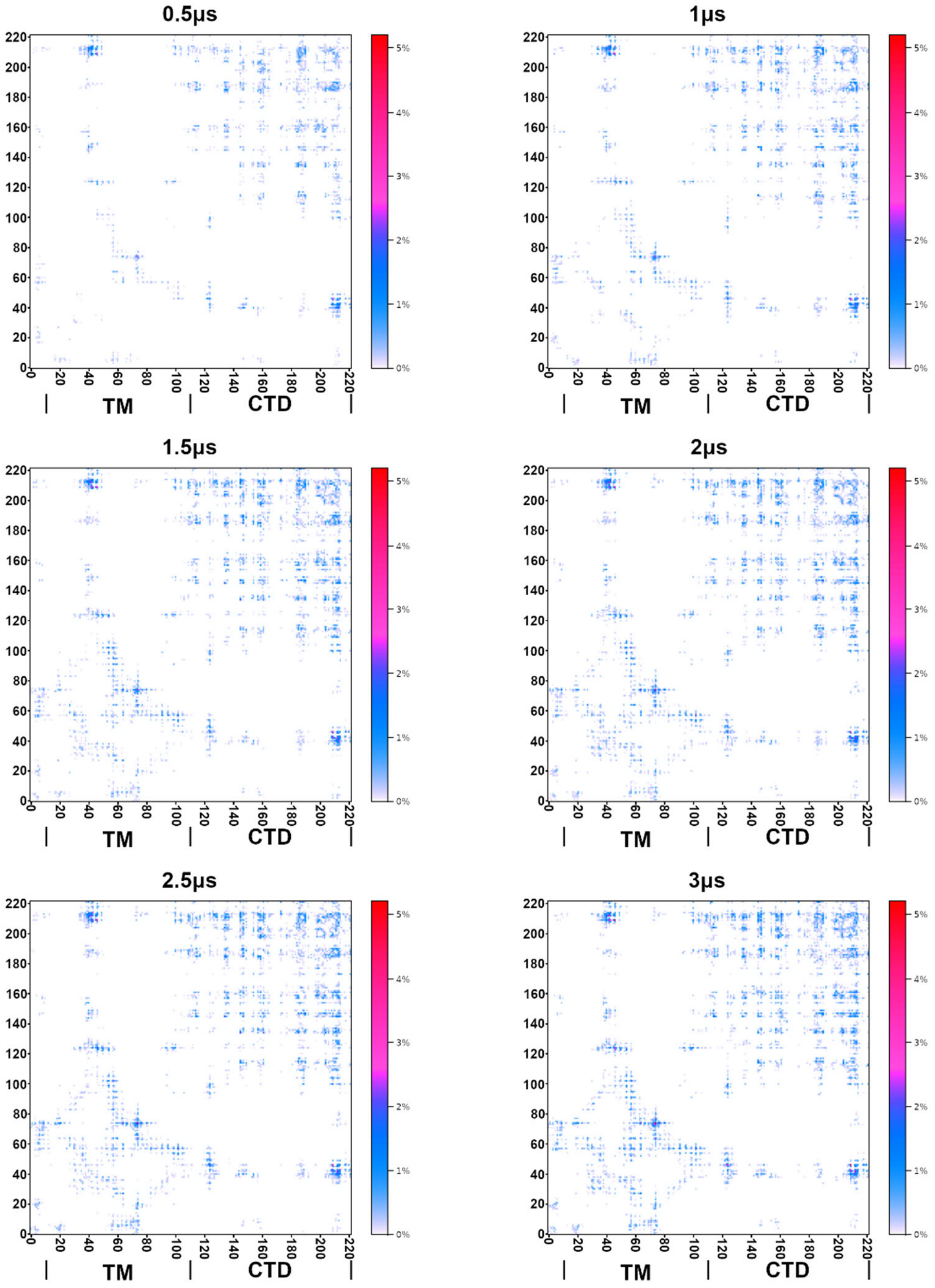


Figure S24. The formation of M dimers. The average contact maps between Ms are shown for several points in simulation time.
